# Seasonal land use land cover change and the drivers in Kafta-Sheraro National Park, Tigray, Ethiopia

**DOI:** 10.1101/2021.09.29.462456

**Authors:** Fitsum Temesgen, Bikila Warkineh, Alemayehu Hailemicael

## Abstract

Kafta-sheraro national park (KSNP) is one of the homes of the African elephant has experienced extensive destruction of woodland following regular land use & land cover change in the past three decades, however, up to date, data and documentation detailing for these changes are not addressed. This study aims to evaluate the land use land cover change and drivers of change that occurred between 1988 and 2018. Landsat 5(TM), Landsat7 (ETM^+^), and Landsat 8 (OLI/TIRs) imagery sensors, field observation, and socio-economic survey data were used. The temporal and spatial Normalized difference vegetation index (NDVI) was calculated and tested the correlation between NDVI and precipitation/temperature. The study computed a kappa coefficient of the dry season (0.90) and wet season (0.845). Continuous decline of woodland (29.38%) and riparian vegetation (47.11%) whereas an increasing trend of shrub-bushland (35.28%), grassland (43.47%), bareland (27.52%), and cultivated land (118.36 km^2^) were showed over thirty years. More results showed bare land was expanded from wet to drier months, while, cultivated land and grazing land increased from dry to wet months. Based on the NDVI result high-moderate vegetation was decreased by 21.47% while sparse & non-vegetation was expanded by 19.8% & 1.7% (36.5 km^2^) respectively. Settlement & agricultural expansion, human-induced fire, firewood collection, gold mining, and charcoal production were the major proximate drivers that negatively affected the park resources. Around KSNP, the local community livelihood depends on farming, expansion of agricultural land is the main driver for woodland dynamics/depletion and this leads to increase resources competition and challenges for the survival of wildlife. Therefore, urgent sustainable conservation of park biodiversity via encouraging community participation in conservation practices and preparing awareness creation programs should be mandatory.

## Introduction

Land use land cover (LULC) change refers to human-dominated modification of the terrestrial surface of the Earth [**1**]. However, the land use/cover pattern of a given region is an outcome of natural and socio-economic factors [**2–4**]. In developing countries land use land cover (LULC) change is an extensively spreading and accelerating process [**5**] which is mainly driven via anthropogenic activities and the change brings significant impact on the terrestrial ecosystem [**6–8**]. Globally, there has been an increasing trend of biodiversity crisis over the last four decades; this is due to the conversion of LULC leading to fragmentation of the natural habitats [**9–12**]. This global loss of biodiversity has great potential to interrupt relevant ecological processes and hinder ecosystem services which are essential for human beings [**13–15**].

Lately, worldwide biodiversity is dominantly concentrated in protected areas (PAs); because the primary objective of establishing protected areas (PAs) is to protect and conserve biodiversity because they have more diversity than other land covers [**16–19**]. However, in Africa, the increasing anthropogenic impacts via fragmentation and loss of wildlife habitats have created an obstacle for the effectiveness of PAs and their value of conservation [**20–22**]. Similarly, in most parts of the world, the activities of neighboring communities of PAs are expected to continue with the potential to negatively influence them [**23, 24**]. Globally habitat conversion is one of the significant issues to threaten biodiversity via landscape cover change [**25**]. LULC change around PAs has direct impacts on PAs biodiversity, ecological process, and habitats health [**26, 27**]. In developing countries, land cover modification of an area around PAs inhibited the effectiveness and capacity of PAs conservation. Over the last three decades, LULC change had been occurring rapidly in PAs and is projected to continue [**21, 28**]. Even in Europe LULC rate of conversion around PAs (~1km buffer zones) is twice inside PAs [**29**]. On the other hand, in Africa, the survival rates of the natural biodiversity around PAs were half of those in PAs [**28**]. As a result of anthropogenic intensification continue evaluating and monitoring of the LULC change in and around PAs are paramount significant [**30**].

The resources in and around PAs are more critical in developing nations of the communities living adjacent to PAs because their livelihoods are often directly dependent on the resources of the land occupied [**31**]. The land use pressure via historical expansion of cultivated land and pasture has largely been at the expense of forests, globally, during the 1990s, there had been an average loss of 16 million hectares of forests per year [**32**]. Agricultural expansion has been reported the main driver for deforestation and consequently loss of terrestrial biodiversity [**33, 34**].To ensure the effectiveness of PAs in developing countries, it is necessary to understand changes driven by the surrounding landscape [**26, 30**]. Thus, monitoring of LULC changes are indispensable aspect for further understanding of change mechanisms and modeling the impact of change on the environment [**35**].

Land use land cover (LULC) change studies have been vastly reported from different corners of Ethiopia in different periods. Most of the studies have documented a considerable expansion of farmland at the expense of other LULC types inside and outside PAs. [**36**] Reported the conversion of forest to agriculture has become a major problem in East Africa due to rapid population growth and subsequent resource competition. In Ethiopia agricultural expansion is the top dominant factor for forest degradation [**37**]. A study in the Eastern Tigray region revealed a strong decrease in the forest and bushland in favor of arable and rangelands [**38**]. Similarly, in the Kafta-Humera district (around the study site) agricultural land has largely expanded by shrinking the coverage of woodland [**39, 40**]. Extensive agricultural expansion as a cost of woodland and dense forest decline was also reported from Nechisar national park [**41**], Bale mountain national Park [**42**], Babile elephant sanctuary [**43**], and Abaya-chamo basin [**44**].

Several studies of LULC change were also reported outside Ethiopia. In Maputaland-Pondoland-Albany biodiversity hotspot (MPA) of South Africa; agricultural land and human settlement showed an increasing trend as a cost of natural forest destruction [**30**]. In similar manner agricultural lands were increased as a conversion of more than three million km^2^ of forest, grassland, and shrublands in PAs of conterminous United States [**45**]. In Sagarmetha national park, Nepal around 26.27 km^2^ natural forests was declined due to an expansion of grazing and settlements [**2**]. Likewise, a study in semi-arid India from 1991 to 2016 about 98% of agricultural land expanded via the conversion of the forest [**46**]. Furthermore, tropical and temperate forest in PAs of Altas Cumbres Mexico was significantly changed to cultivation [**47**].

Land use land cover (LULC) change is key information for scholars who are working in land management studies [**48**]. Therefore, understanding the dynamics and driving forces LULC changes at the local and global levels is fundamental to develop strategic planning and the analysis of land-related policies [**49**]. To announce each land cover change, remotely sensed (RS) and Geographical information systems (GIS) are widely used data sources [**48, 50**]. RS multitemporal satellite imagery plays a vital role in quantifying spatial and temporal phenomena and understanding the landscape dynamics [**7, 51**]. Remotely sensed (RS) data and techniques use globally available satellite imagery, making it a widely accessible research methodology for scientists in both developed and developing nations [**30**]. These techniques made it possible to evaluate LULC change at a low cost [**5**], in less time couple with reasonable accuracy [**51**]. Comparatively, the combination of RS data and field observations can accomplish LULC classification, change detection, and drivers of change, more accurately and acceptance than separately [**20, 52**] because satellite data analysis alone can miss the drivers of LULC change [**53**]. Moreover, LULC change analysis acceptance and benefits were maximized when satellite image analysis mixed with socio-economic (local residents) participation [**2, 42, 54, 55**].

Worldwide particularly in the developing nations the most common and free charge accessible satellite sensors applied for LULC dynamics detection and evaluation are Landsats namely; Thematic mapper (TM), Enhanced thematic mapper plus (ETM^+^), and Operational land imager (OLI) [**56,57**]. These Landsat imagery sensors have the capacity of ease of access and temporal coverage to analyze land-cover change [**47**]. Therefore, for this study Landsat-5 of 1988 and 1998, Landsat-7 of 2008, and Landsat-8 of 2018 sensors were applied. Thus, the main objectives of this study were (1) to assess the pattern of land use land cover change in Kafta-sheraro national park (KSNP) for three decades (1988 to 2018) (2), to compare the seasonal variation of LULC change, and (3) to evaluate the drivers of LULC change and its effect on conservation of the park.

## Materials and methods

### Description of the study area

Kafta-Sheraro National Park (KSNP) was designated as a park in 2007 (Letter, N<u>o</u>: 13/37/82/611) with an area of 2176.43 km^2^. While the park was formerly named “Shire wildlife reserve” was established in 1973 with an estimated area of 750 km^2^ governed by Tigray national regional state. Kafta-Shirero national park (KSNP) is located in Kafta-humera and Tahtay-adyabo weredas (districts) of Western and Northwestern Zones of Tigray region 1356 km far from Addis Ababa and 490 km of Mekelle city. The park is situated in the North of Ethiopia between latitude 14°05′-14° 27′ N and longitude 36°42′-37°39′ E. The park is bordered by Eritrea in the north and transverse by the Tekeze River (Fig 1). The elevation of the Park varies from 539 to 1130 meters above sea level (m.a.s.l). The landforms of the areas are heterogeneous in nature and consist of a flat plain, undulating to rolling; some isolated hills and ridges, a chain of mountains, and valleys [**58**].

**Fig 1:**
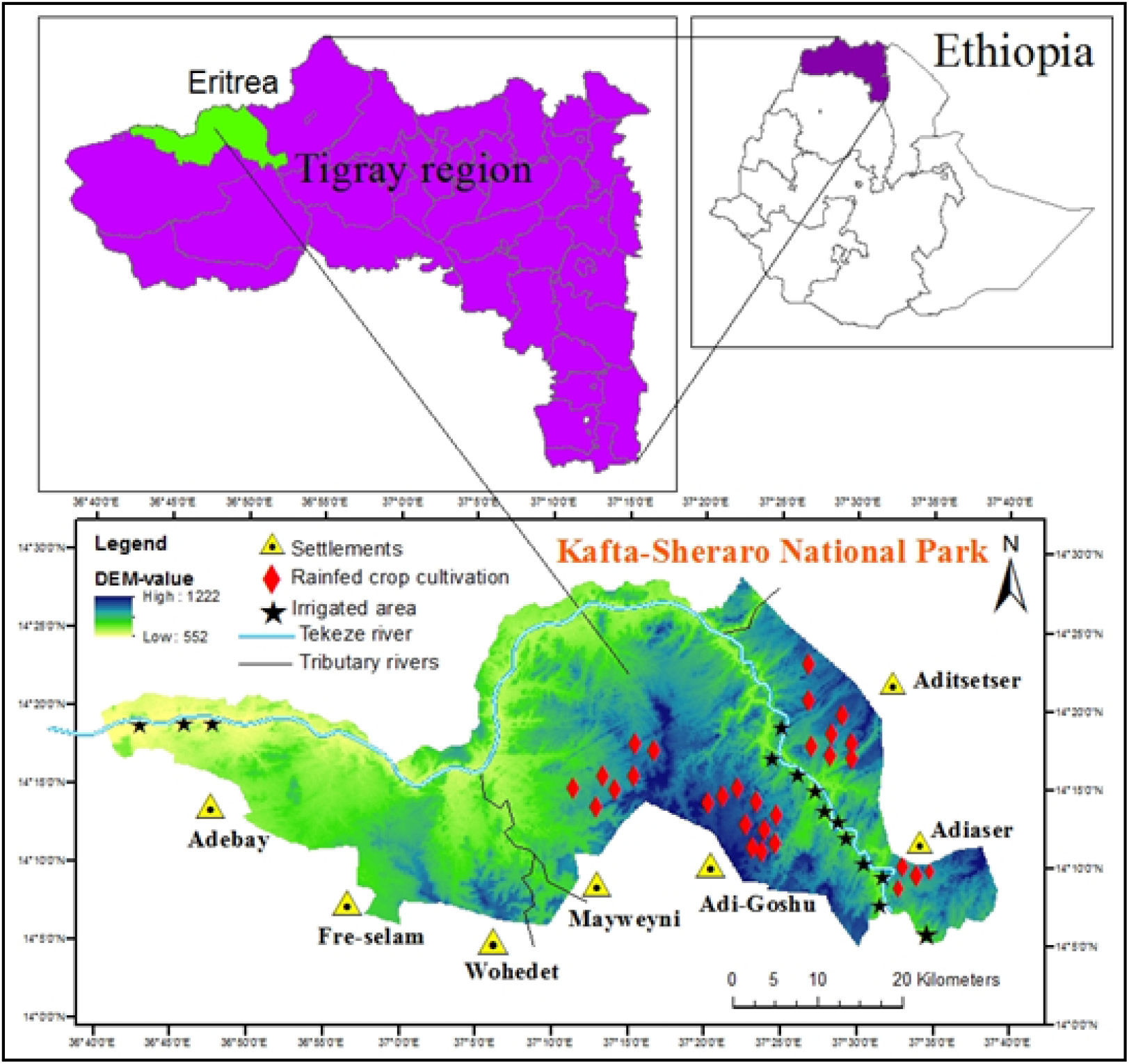
Location map of the study area

The climate of the area is generally characterized by hot to warm semi-arid and seasonal rainfall [**58**]. The maximum monthly temperature is in April (43.7 °C) while the minimum monthly is in December (19.2 °C) and January (19.1 °C) respectively. The mean monthly temperature ranges from 28.35 °C to 35.1 °C. The coolest temperature occurs in August while the warmest temperature occurs from March to May. The rainfall pattern is greatly varied with months of the season. The short rains occur in June and September and the long rains occur during July (174 mm) and August (252 mm) whereas rare cases of rains in the rest months appeared (Fig 2).

**Fig 2:**
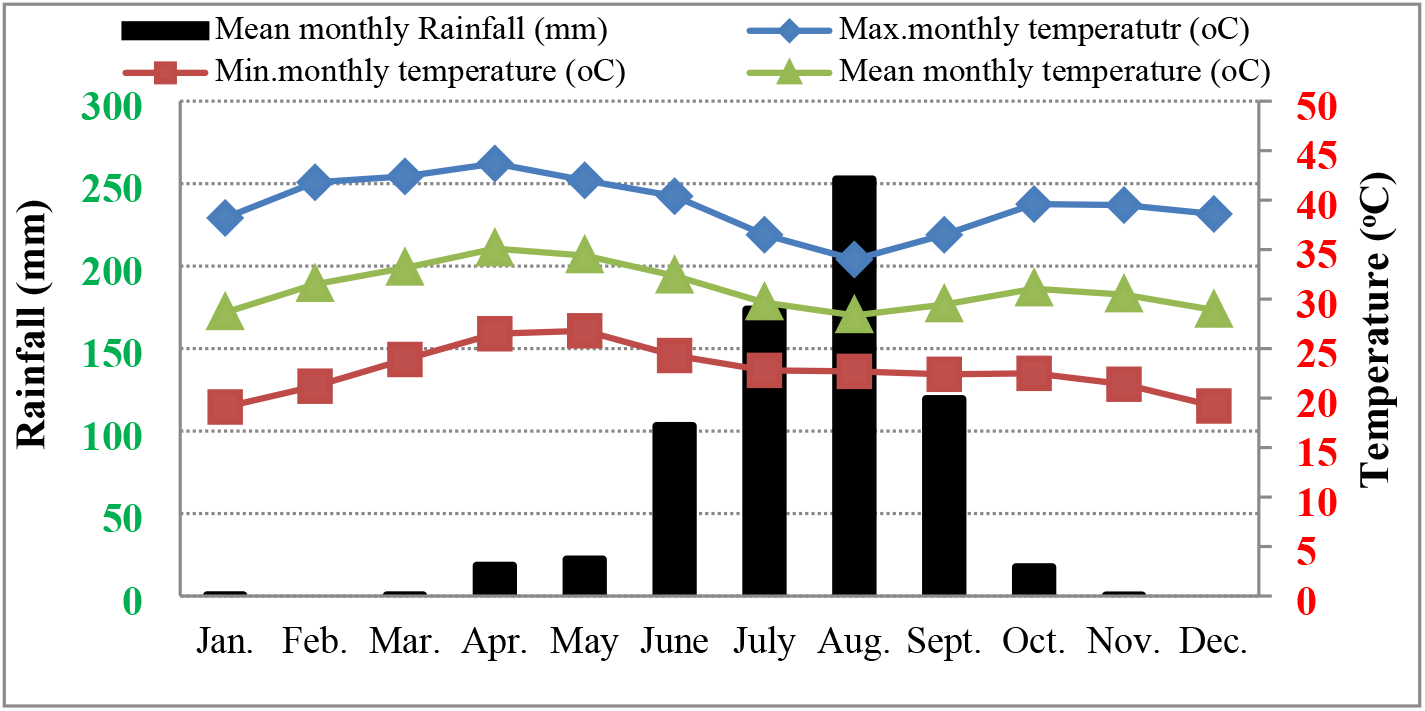
Maximum, mean, and minimum monthly Temperature (°C) and mean monthly Rainfall (mm. month^-1^) of Humera and Shiraro districts Meteorological Center from 1996-2016 [**60**].

The KSNP harboring above 70 woody species, 46 trees, 18 shrubs, and 6 tree /shrub. The most dominant and frequent tree species of the park are *Acacia mellifera, Combretum hartmannianum, Terminalia brownii, Balanites aegyptiaca, Dicrostachy scinerea, Acacia senegal, Acacia oerfota, Boswellia papyrifera, Ziziphus spina-christi*, and *Anogeissus leiocarpus* (**58**). The park is also home to large mammals like African elephant, Roan antelope, Oribi, Spotted hyena, Greater kudu, warthog, Anubis baboon, Grivet monkey, crocodile, fish species and wintering migratory bird (Demoiselle crane) along the Tekeze River (Field observation).

Agriculture is the main source of livelihood and economic activities of the study settlers. The livelihood of the local communities of the district (i.e Kafta-humera wereda surrounding the park) is dominated by mixed farming of crop-livestock production [**59**].

### Data collection and sources

#### Satellite image data

In this study, Landsat 5 thematic mapper (TM); Landsat 7 Enhanced thematic mapper plus (ETM^+^), and Landsat 8 Operational land imager/Thermal infrared sensor (OLI/TIRS) multi-spectral satellite sensors data were used to detect LULC change between 1988 and 2018 (Fig 3). The images were downloaded from the Earth Explorer (http://earthexplorer.usgs.gov) and covering a time span of 30 years period. The park is covered by Landsat frame path/row (170/50) of the Worldwide Reference System. A detail explanation of each satellite sensor is described in Table 1. The images with high resolution and minimum or no cloud cover were selected from a number of images for each period to minimize errors or confusion for LULC classification. A total of 26 images for the dry and wet season, 5 for LULC change detection, and 21 for normalized difference vegetation index (NDVI) analysis were downloaded. In this study, the dry season period was defined from 24 October to May and wet season was defined from June to 23 October. The images for the wet season classification were taken between September and 23 October while for the dry season between February and April. Consequently, these months were preferred for all satellite image sensors because they were found to have no or minimum cloud cover and water vapors.

**Fig 3:**
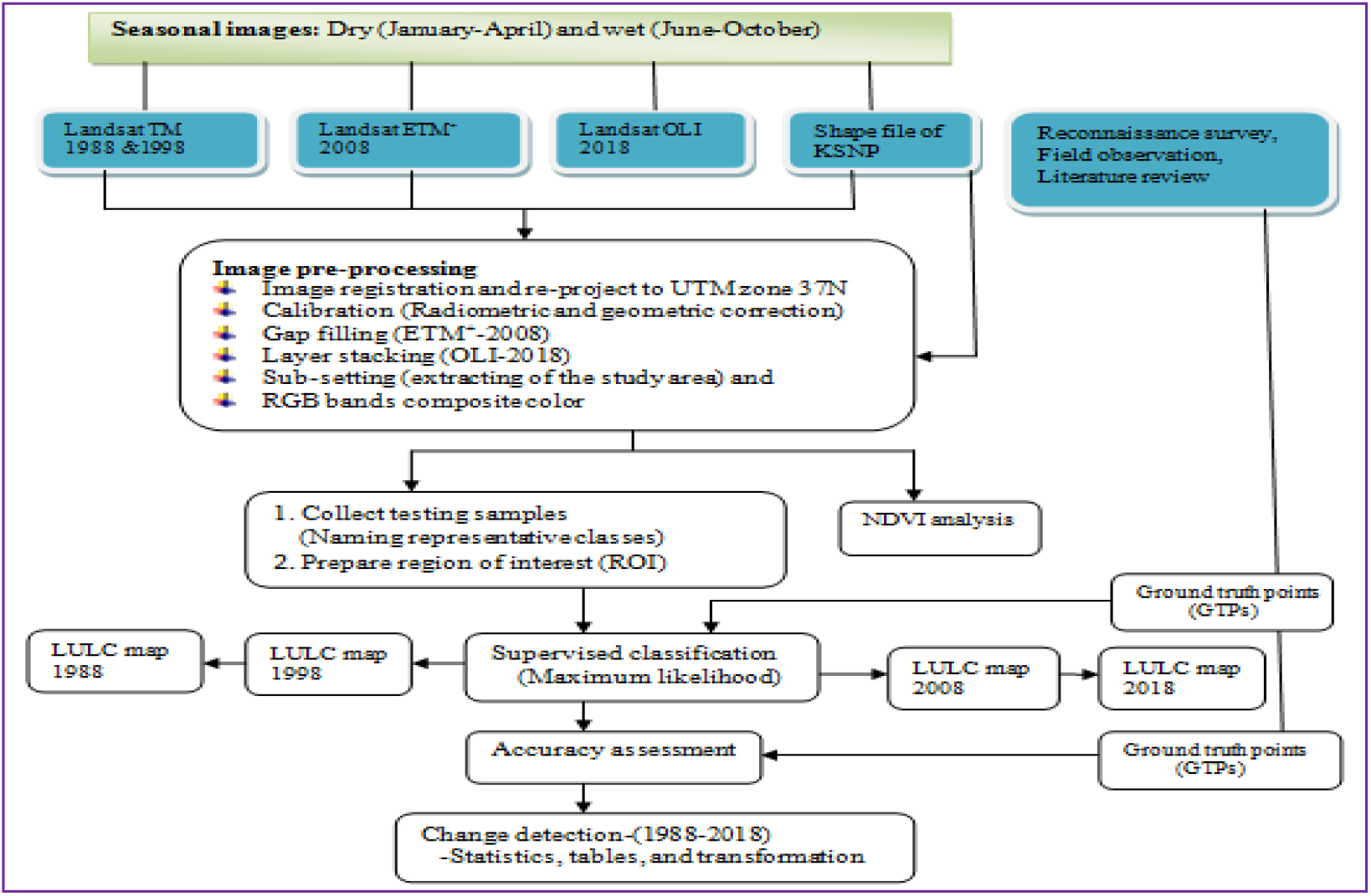
The flow diagram summary of steps and data utilized in image analysis

**Table 1:** Data type and detail description of satellite images used in LULC change analysis

| Satellite /Sensor type | Path Row | Acquisition date | Spatial Resolution (m) | Bands (B) used for spectral signature | Bands wave length (μm) |
| --- | --- | --- | --- | --- | --- |
| Landsat-5 TM | 170/50 | 23-10-1988 | 30 | B1,B2,B3,B4,B5,B7 | 0.48-2.20 |
| Landsat-5 TM | 170/50 | 19-10-1998 | 30 | B1,B2,B3,B4,B5,B7 | 0.48-2.20 |
| Landsat-7 ETM<sup>+</sup> | 170/50 | 06-10-2008 | 30 | B1,B2,B3,B4,B5,B7 | 0.48-2.22 |
| Landsat-8 OLI/ TIRS | 170/50 | 10-10-2018<br>16-03-2018 | 30 | B2,B3,B4,B5,B6,B7 | 0.45-2.29 |

TM= thematic mapper, ETM<sup>+</sup> = enhanced thematic mapper plus, OLI= operational land imager, TIRS= thermal infrared sensor and B=band, and calander order (day-month-year)

### Field observation data

Field visits were carried out for identification of major LULC types and to take training points that are changed seasonally in the Kafta-Sheraro National Park (KSNP). The survey took from December 2018 to April 2018 for a dry season while from mid-June 2018 to begining of November 2018 for wet season classification. Table 2 shows each season change of LULC identified in the field and cross-checked through interview of focus group discussion. Moreover, field training support by the local farmers having farmland inside and around the park. Accordingly, the local farmers’ interviewed that the plowing and sowing time of rainfed crops starts from June and the crops are harvested from mid-November to December. For example, after the rainfed crops are harvested they could leave the land bare until the next sowing year (June) or left totally free and shifted to a new area. Similarly, the grasses cover in the wet season appeared bare land until the next rain season (Table 2).

**Table 2:** Some of the LULC classes change seasonally or remain the same and the field point’s seted for the two seasons (wet and dry)

| Land use land cover classes | Wet season | Dry season |
| --- | --- | --- |
| 1. Rainfed crops | Cultivation | Bare land |
| 2. Grazing/grass cover | Grass land | Bare land |
| 3. Irrigated land | Cultivation | Cultivation |
| 4. Streams & main tributary rivers | Water body | Bare land |
| 5. Main road | Bareland | Bareland |
| 6. Sandy and small gravel area | Bareland | Bareland |

Fieldwork is mainly focused on observing and capturing the various LULC using a digital camera and each sampling location was recorded via geographical positioning system (GPS) of handheld GARMIN GPS-60.To emphasis, the classification more accurately; above 100 ground truth (latitude and longitude record) were collected from each LULC type of 2018 image and a total of above 700 points from seven classes for classificaion & accuracy assessment. The accuracy assessment was basically well done for the Landsat-8 (OLI) of the 2018 satellite image because the points directly show the recent feature of LULC categories. The ground control points were divided into two groups, one group for selecting training sites for the supervised classification and the second group for the accuracy assessment.

### Socio-economic survey

#### Sampling design

The study has collected the drivers of LULC change and related issues from three basic sources namely: (1) semi-structured household questionnaires, (2) focus group discussions, and (3) key informant interviews. Different sampling procedures were used to select the sampled Kebeles and households. As the Park is located in two weredas (districts) of Kafta-humera and Tahtay-adyabo, seven Kebeles (the smallest administrative units of Ethiopia) were purposively selected from the total, based on proximity to the park and their livelihood directly dependent on the resources of Kafta-sheraro national park (KSNP) and its surrounding area. A systematic random sampling method was used in order to select the representative sample respondents for the household interviews from individual kebeles whereas the purposive sampling technique for focus group discussion and key informant interviews. The sample size for the present study was calculated using equation (1) sampling technique [**61**].

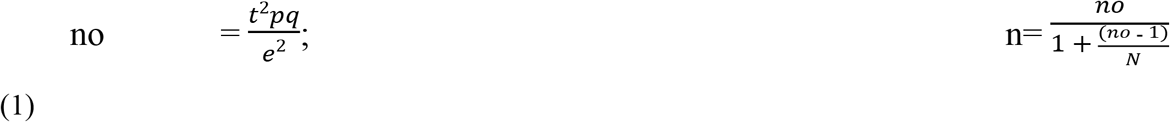

**Where,** no= assumed simple random sample size of house holds (384); p=estimated proportion of the population to be included in the sample (i.e. 50%); q=1-p; t= uncertainity (number of standard error) in the number of people depend on park resources of ±5% (at 95% confidence interval level, Z-value =1.96); e= the margin of error (0.05); n=sample size, and N=the total number of house hold heads (i.e 5458).Using equ.1 the result of sample size was 359; however, to compensate and cover the non response of house holds, the sample size was increased by 10% (36). Therefore, the total sample size interviewed in this study was 395 (Table 3).

#### Household survey

The questionnaires had both open and closed-ended questions to gather information about the perceptions of the local communities on LULC changes and the drivers of change in KSNP between 1988 and 2018. The questionnaires have covered 395 households during November 2018-2019 from seven Kebeles (Table 3). For individual household response took 50-70 minutes. The questionnaires also include local people particularly elders were consulted about the age and history of the land use type and identified the main drivers of LULC change of KSNP using open-ended questions, that is the periods in which land use/cover type remained in the same state or not. The targeted populations for semi-structured interviews were park nearby community (villagers) having direct interaction with KSNP, irrigation farm holders’, and seasonal livestock owner. The questionnaires had basically designed to gather general household characteristics, forest coverage trends, perception of the local people on LULC change, and the drivers of change (Appendix S1). Finally ranked the illegal activity for cover change based on their degree of impact on the sustainability of the park.

**Table 3:** Total selected sample household heads from seven kebeles of Kafta-humera and Tahtay-adyabo districts (Source: survey 2018)

| Wereda (district) | Selected Kebeles | Total heads of house hold | Sample size has taken from each kebele |  |  |
| --- | --- | --- | --- | --- | --- |
|  |  |  | Male | Female | Total |
| Kafta-humera | Adebay | 2976 | 88 | 36 | 124 |
|  | Fre-Selam | 291 | 25 | 9 | 34 |
|  | Wohedet | 262 | 21 | 7 | 28 |
|  | May-Weyni | 265 | 22 | 8 | 30 |
|  | Adi-Goshu | 671 | 55 | 17 | 72 |
| Tahtay-adyabo | Adi-Aser | 324 | 28 | 9 | 37 |
|  | Adi-Tsetser | 669 | 51 | 19 | 70 |
| Total |  | 5,458 | 293 | 102 | 395 |

Focus group discussion (FGD): this is away of data collection which involves the investigator gathering a group of participants together to discuss relevant issue of the study. The FGD was performed within seven Kebeles (villages town) of the study area by involving the elder people who are more knowledgeable in discussions and older than 60 years and who were lived in the area for more than 28 years. This has helped us to aware more about the ongoing LULC change (extent, trend, and past and present drivers of LULC change) in the study area. We took a total of seven focus group discussion within each group had 6-10 persons as limited by [**65**] and the duration for each discussion were 120-140 minutes.

#### Key informant’s interviews (KII)

According to [**66**] report key informant interviews are qualitative in-depth interviews that involve the researchers in addressing the issue to the people who have knowledge about what is happening in the community. The qualitative data collection from key informants was via direct individual interviews and focus group discussions. In our study, the main objective of key informant interview was to collect detail information from a specific group of people like community leaders, elderly group and professionals who have firsthand knowledge about the ongoing problems happened in KSNP by the communities. Thus, the researcher has conducted key informant’s interviews with experts of crop production, livestock production, forest and wildlife conservation, soil and water conservation of the districts, and the staff members of the park.

### Data analysis

#### Pre-processing of image

Before LULC classification and detection of changes pre-processing of satellite images is an imperative process with the aim of developing an inline association between the biophysical phenomena on the ground and the acquired data [**67**]. Before all activity, Landsat images (TM 1988 and 1998, ETM^+^ 2008 and OLI 2018 imageries) were geometrically rectified (geocoded) to the World Geodetic System 1984 (WGS 84) and set a projection to Universal Traverse Mercator (UTM) zone 37N specific to Ethiopia. Geometric and radiometric (reflectance) calibration: during image acquisition satellite images have different types of distortions/noise and this reduced the quality of the image. Thus for better performance of the Landsat time series of LULC change analysis, consistent image sets of geometric and radiometric corrections are the two significant activity [**68,69**]. The calibration of landsat imagery was performed based on the known solar geometry and on the gain and bias values provided by the Landsat metadata [**70**]. For the present study geometric and radiometric (reflectance) corrections were carried out to decrease negative atmospheric effect or correct for changes that occurred in scene illumination, atmospheric solar condition, and viewing geometry as applied by [**71**]. Likewise, all images having cloud and cloud shadow were removed using cloud mask (*fmask function*) under ArcMap10.5 tool of image analysis. Other additional pre-request activities undertaken following calibration were gap filling, layer stacking, and sub-setting of bands. Color combination of bands: refines image interpretability via increasing differentiability among objects of the image for classification. Basically, Red, green, blue (RGB) color composites arrangement is common in any satellite image display. In this study for better visualization of different objects of the images, we created color combination by taking band 7 for infrared (2.064-2.345 μm), 4 for nearinfrared (1.547-1.749 μm), and 1 for blue (0.772-0.898 μm) were chosen for the images TM 1988, 1998 and ETM^+^ 2008. As for the image OLI 2018, the band 7 for infrared (2.107-2.294 μm), 5 for near-infrared (0.851-0.879 μm), and 4 for red (0.636 to 0.673 μm) were chosen. All the image pre-processing activities have been done using ENVI 5.3 software.

#### Land use/cover classes classification

The images were classified using the supervised classification algorithms under ENVI 5.3 because we are familiar with the study landscape. This classification method may be more preferable for LULC change detection if prior information about the landscape is gained through personal knowledge of the study area [**72**]. The individual LULC class signatures of polygons (training areas) were marked based on the field observation, household knowledge, and color combination of bands (image visual interpretation). Then the image data set in LULC class is placed via Maximum Likelihood Classifier (MLC). Even though there are different classifiers, the MLC algorithm was more performed using all the spectral bands fit to vegetation. MLC was reported the most common, successful, and widely adopted classification algorithm [**3, 73–76**]. This technique has also a greater probability to weight minority class that can be swamped by the large class during training samples taken from images. Because the minority classes in the image have the opportunity to be included in their respective spectral classes (reduce uncategorized pixel) from entering into another class [**77**]. Accordingly, seven major LULC classes were recognized in KSNP. Therefore, the description of the LULC classes was based on the author’s prior knowledge of the study site, detailed, and consecutive field observations (Table 4).

**Table 4:** Land use land cover catagories and their explanations in Kafta-sheraro national park

| LULC type | Description |
| --- | --- |
| Woodlands | Large and medium trees which have medium canopy cover arise from lower range of grasses and mixed herbs and grasses. |
| Shrub-bushlands | Dominantly covered by short height shrub bush structure of plants and arise from mixed lower coverage of grasses and herbs. |
| Riparian forests | Dense canopy cover vegetation of the park along Tekeze River and its tributary rivers valley having vertical stratification and dominated by tall trees or wood lands. |
| Grasslands | Plains, rough ground and relatively hilly areas cover by different predominantly grass species mixed with herbs and natural pasture for animals. |
| Agricultural land | Both rainfed crops and irrigated vegetable and fruit crops like banana and mango plantation. Areas of store house and farmhouses are also under agricultural land. |
| Water bodies | Water courses of Tekeze river, permanent & seasonal tributary rivers, and streams |
| Bareland | Areas including no vegetation cover, gravel and asphalt roads, degraded lands, bare ground and gold excavation areas, mixed sand and small gravel which are found in Tekeze River sides and its tributary rivers. |

### Accuracy assessment

Accuracy assessment is useful to assess the quality of the data collected in the field and the classified images. This technique determines the sources of error encounter during the classification of satellite images [**78**]. We compare the accuracy assessment for dry and wet seasons of 2018 satellite imageries. Accuracy assessment determines how accurate the referenced or ground-truth data region of interest agreed with classified images of the remotely sensed data in which precision testing was conducted using the Kappa index [**79–81**]. Accuracy assessment was expressed by the following four parameters from equations 2-5 [**78, 82, 83**].

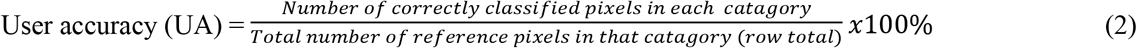

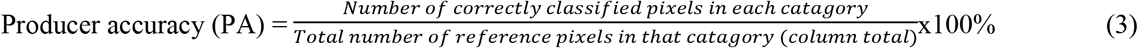

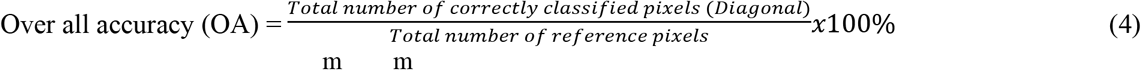

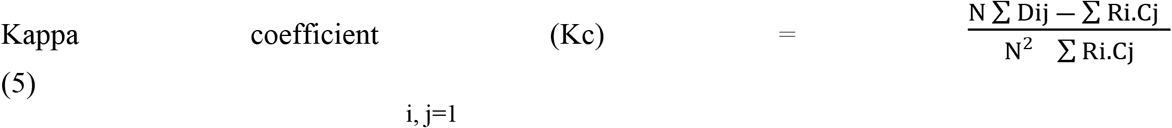

Where, N =total number of pixels, m = number of classes, Σ Dij=total diagonal elements of an error matrix (the sum of correctly classified pixels in all images), Ri =total number of pixels in row i, and Cj=total number of pixels in column j.

### Landuse landcover change analysis

The magnitude of change is a degree of expansion or reduction in the LULC size of the classes. A negative value presents a decrease in LULC size while a positive value indicates an increase in the size of LULC class [**84**].The percent rate of LULC change between periods were computed following equation (6) [**46, 85–87**].

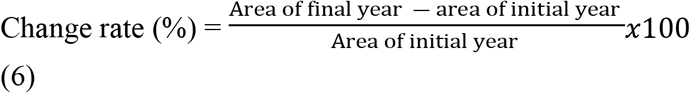

The annual rate of change per year was calculated using equations (7) and (8) [**88, 89**].

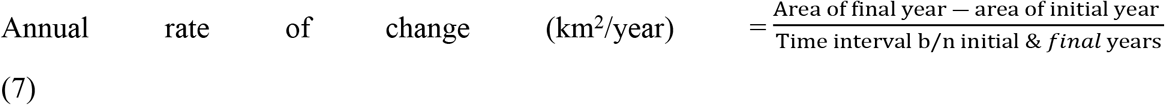

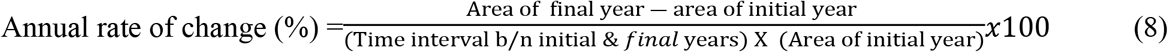

### Normalized difference vegetation index (NDVI) analysis

Normalized difference vegetation index (NDVI) evaluates the relationship between spectrum reflectance variability and change in vegetation growth rate [**90**]. It is used to distinguish the status of vegetation cover, state of vegetation degradation, and the extent of forest loss [**91**]. As the study area is found in the dryland region and anthropogenic activity is also relatively high, NDVI information is relevant to assess the vegetation cover status. In our study Landsat imageries of 1988 and 2018 were utilized to extract NDVI values of vegetation cover change classification: Non-vegetation, sparse vegetation, and high-moderate density vegetation (Fig 7 and Table 9). The NDVI value ranges between −1 and +1. As the value increases towards +1 or increasing positive NDVI values indicate the dense vegetation (vegetated the plant canopy), and close to zero or decreasing negative values (−1) indicates the non-vegetation surface such as water and bare ground [**92**]. A high positive value of NDVI is also computed from vegetated agricultural cover crops [**93**]. NDVI is calculated based on the difference in the ratio of Red (R) and near-infrared (NIR) reflectance, equations (9) and (10) [**90, 93, 94**].

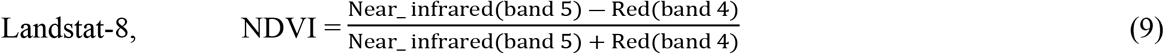

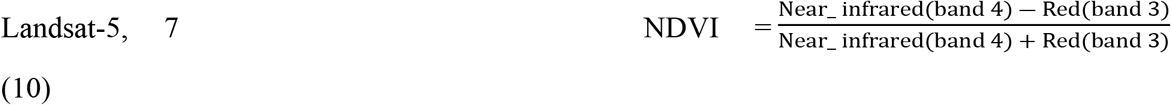

**<u>Note</u>:** in this analysis the index was computed based on the difference in Red band 4 (0.64-0.67 μm) reflectance and NIR band 5(0.85-0.88 μm) reflectance of Landsat-8 OLI of both dry (March-2018) and wet season (October-2018) satellite images. In addition for the TM and ETM^+^ sensors Red band 3(0.63-0.69 μm) and NIR band 4(0.77-0.9 μm) reflectance.

The study site is found in semi-arid region, where climate variables are limiting factors for vegetation cover determination. From the climate variables precipitation has a direct relation on spatial and temporal change of NDVI. Due to the absence (discontinuous) of remotely sensed satellite images between 1996 and 2006, the NDVI relation with climate variables analysis was conducted between 2007 and 2016. Because in these periods,a continuous satellite image data were available directly matched with the recorded precipitation and temperature of the same years and seasons. Thus, the statistical relationship between NDVI response and precipitation and/or temperature separately was examined through linear regression model as applied with the equation (11) for one decade data.

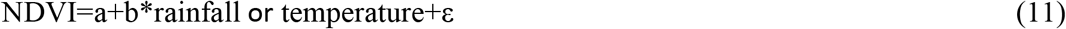

**Where**,’NDVI’ =dependent factor; rainfall or temperature=independent factor, a=intercept, b, =partial slope coefficient for variable rainfall or temperature, and *ε*=the random error. The seasonal mean NDVI, annual seasonal rainfall, and seasonal mean temperature were analyzed for the period of 2007 to 2016.The significance mean NDVI change was evaluated at 95% confidence level (p<0.05).

### Socio-economic and associated statistical analysis

The data collected from the seven kebeles (districts) of sampled households were quantitatively analyzed whereas data from focus group discussion (FGDs) and key informant interviews were analyzed qualitatively. Primary, all the socio-economic data derived from the questionnaire were entered into Microsoft excel and arranged as suitable for analysis. Descriptive-statistics analysis (mean and percentage) was used to describe socio-economic variables of the households and summarized their responses using tables and figures. Statistically, the interviewees’ awareness of the drivers of LULC change response between selected socio-economic variables was analyzed by Pearson’s Chi-square (non-parametric test). The main drivers of LULC change obtained from the household surveys were summarized by the ranking method. In this method, the index was calculated as a ranking ratio with the principle of weighted average following the mathematical formula adopted by [**95**] as described in equation (12).

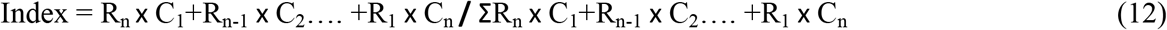

**Where,** Rn =the value given for the least ranked level by respondents (in this study the least rank is 4^th^, then R_n_ = 4, R_n-1_ = 3, R_1_ = 1); C_1_ =the counts of the 1^st^ ranked, C_2_= the counts of the 2^nd^ ranked… & C_n_ = the counts of the least ranked level (in this study, C_n_ = the count of the 4^th^ rank) On the other hand, the local people’s awareness status on LULC change drivers of KSNP which is determined by the socio-economic descriptive variable difference of the sampled households was tested by logistic regression analysis. Logistic regression analysis is the interaction among the driver data and the LULC change category. [**96**] Stated logistic regression is useful for the discrete dependent variable which has a binary (bivariate) outcome and investigates the association of dependent (response) variables with independent (predictor) variables. Therefore, the interlinked mathematical formula is summarized in equation (13).

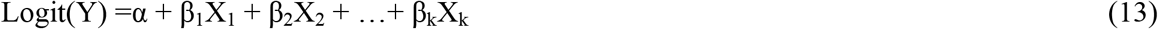

**Where,** Y=the dependent (response) variable; α =the intercept; β_1_ … β_k_ =the coefficients of explanatory variables, and X_1_ … X_k_= the independent (explanatory) variables.

In the present study, the response variable was household awareness difference on the drivers of LULC changes, while independent variables held sampled household descriptive information. Hence, there were seven determinant socio-economic variables used in the analysis include: age category, gender, household size, education level, settlement duration, agricultural land size, and distance from settlement to KSNP border. Thus, this analysis evaluated the impact probability of the independent variables on the dependent variables. Additionally, the correlations between the trends of different LULC change types were computed and a statistically significant association has performed a p<0.05. All the listed above statistical tests (i.e Pearson’s Chi-square (*X^2^*), regression and correlation) were undergone through R-statistical Package [**97**] and IBM SPSS statistics [**98**].

### Ethics approval and research site permission

The study **Ethics** was reviewed and approved by College of Natural & Computational Science, Research Department Graduate Committee (DGC, 2018) & Ethiopian wildlife conservation authority Research Ethics Review Committee. Prior to field visit & data collection, the **permission letter** was gained from Ethiopian wildlife conservation authority for the selected study site of Kafta Sheraro National Park (KSNP). Before distributed the questionnaire the objective of the study was briefly explained to participants & verbal consent was collected. The questionnaire excluded personal identifiers (privacy) like name of interviewee.

## Results

### Landcover classification map and accuracy assessment

Wet and dry season land use land cover (LULC) products assessment revealed that the map accuracies from the maximum likelihood supervised classification approach were good. Based on the ground truth recorded data of 2018 the overall accuracies for images of wet and dry seasons were 86.9% and 91.96% with Kappa coefficients of 0.845 and 0.90 respectively (Tables 5 & 6). The user’s accuracies (UA) and producer’s accuracies (PA) of the wet season woodland, riparian forest, agriculture, and water body were reasonably classified, both the UA and PA of greater than 88% for the tested year. Shrub-bushland of UA accounts above 88%; however, PA value is 77.31% (Table 5). Dry season riparian forest, irrigated land, river water, and bare land were relatively classified well, above 91% for all tested maps. However, woodland and shrub-bushland cover relatively poorly classified with UA ranged from 76.47% to 92.94% and PA 72.48% to 97.5% (Table 6). Fig 4 shows the wet season spatial distribution map of KSNP for the year 1988-2018 and Fig 5 shows the seven dry season land cover classes derived from the 2018 Landsat image.

**Table 5:** Wet season confusion matrix for 2018 OLI classified image

| Classified data | Ground truth data |  |  |  |  |  |  |  | UA (%) |
| --- | --- | --- | --- | --- | --- | --- | --- | --- | --- |
|  | Wood land | Shrub bushland | Riparian forest | Grass land | Agricultural land | Water body | Bare land | Total |  |
| Woodland | 129 | 0 | 7 | 0 | 1 | 0 | 0 | 137 | 88.97 |
| Shrub-bushland | 6 | 92 | 0 | 6 | 0 | 0 | 0 | 104 | 88.46 |
| Riparian forest | 0 | 0 | 128 | 0 | 0 | 0 | 0 | 128 | 96.80 |
| Grass land | 0 | 22 | 0 | 146 | 1 | 0 | 14 | 183 | 73.00 |
| Agricultural land | 0 | 0 | 0 | 1 | 88 | 0 | 0 | 89 | 98.88 |
| Water body | 1 | 0 | 0 | 0 | 0 | 74 | 0 | 75 | 98.67 |
| Bareland | 0 | 5 | 0 | 18 | 0 | 0 | 88 | 111 | 79.28 |
| Total | 136 | 119 | 135 | 171 | 90 | 74 | 102 | 827 |  |
| PA (%) | 88.36 | 77.31 | 89.63 | 82.49 | 93.62 | 98.67 | 85.4 |  |  |
| Over all accuracy | 86.9% |  |  |  |  |  |  |  |  |
| Kappa coefficient | 0.845 |  |  |  |  |  |  |  |  |

Note: classes are shown by no of classified pixels, UA=user accuracy, PA=producer accuracy

**Table 6:**
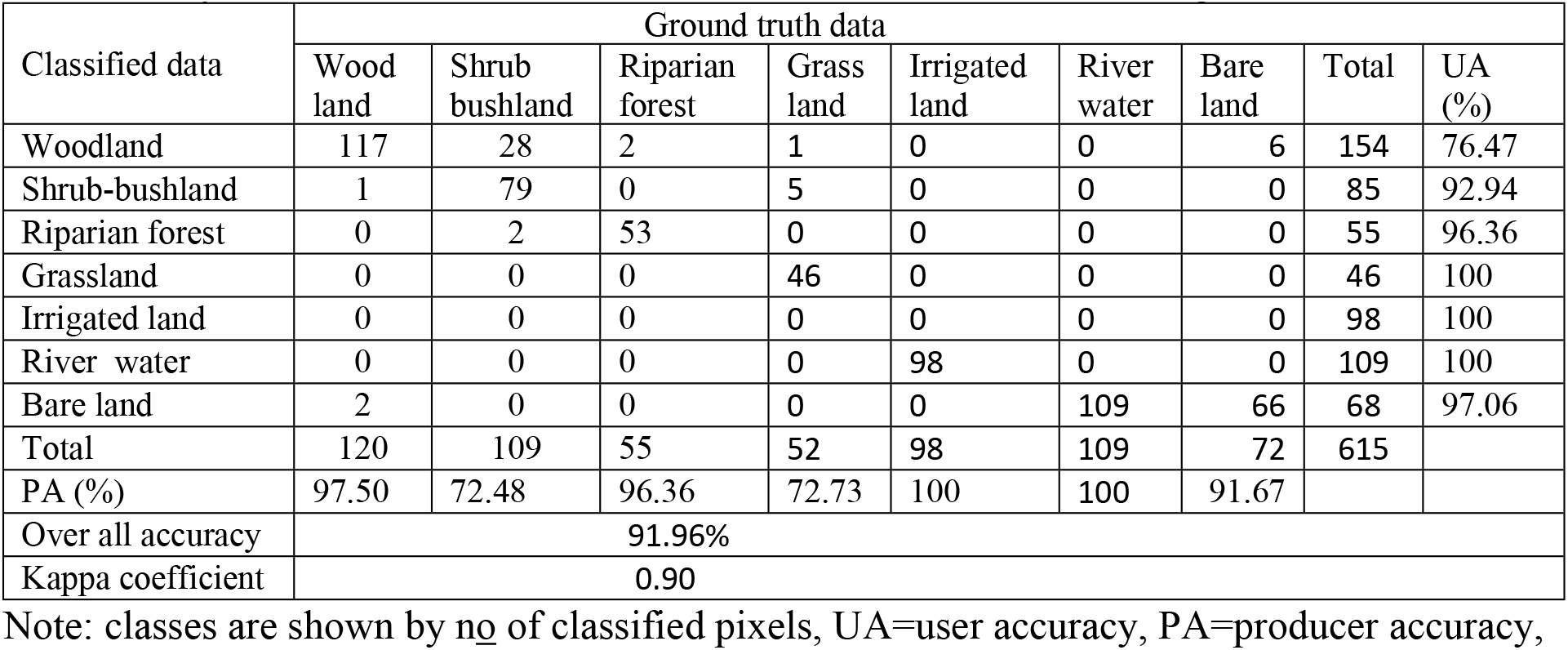
Dry season confusion matrix result for 2018 OLI classified image

**Fig 4:**
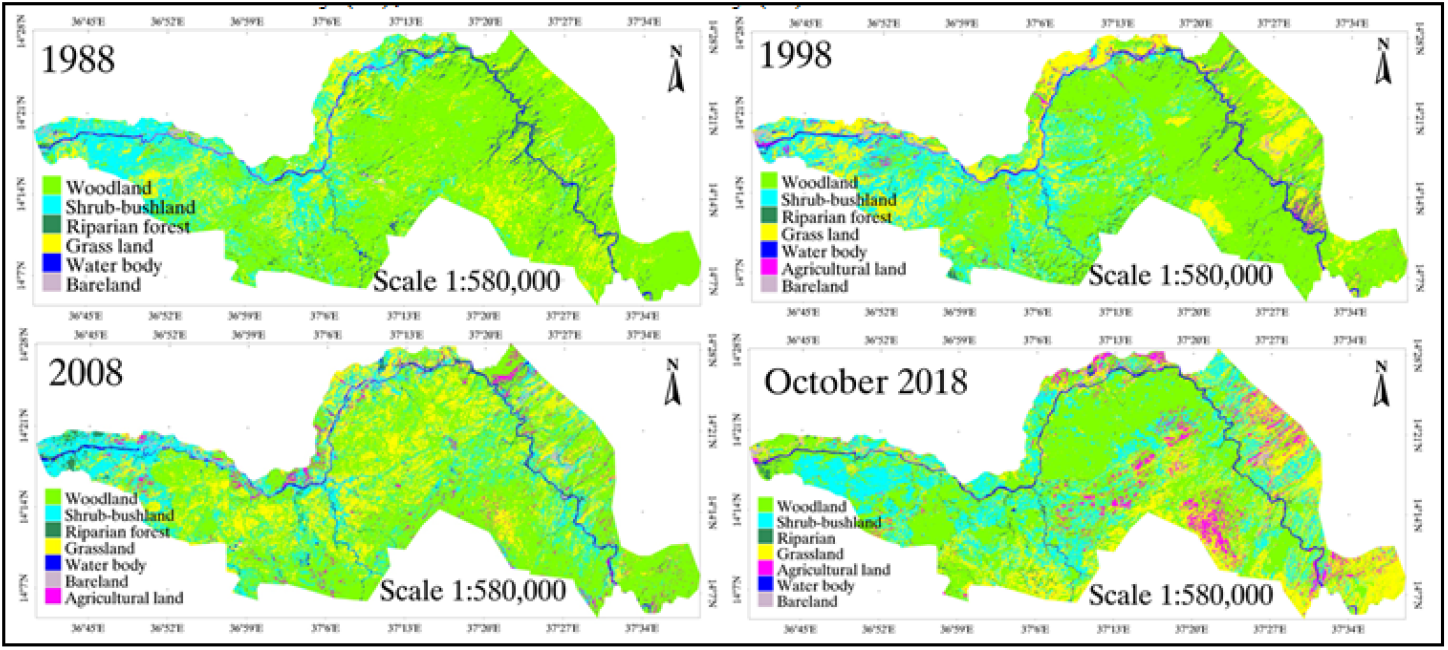
Wet season LULC classes map derived from 1988, 1998, 2008, and 2018 Landsat images

**Fig 5:**
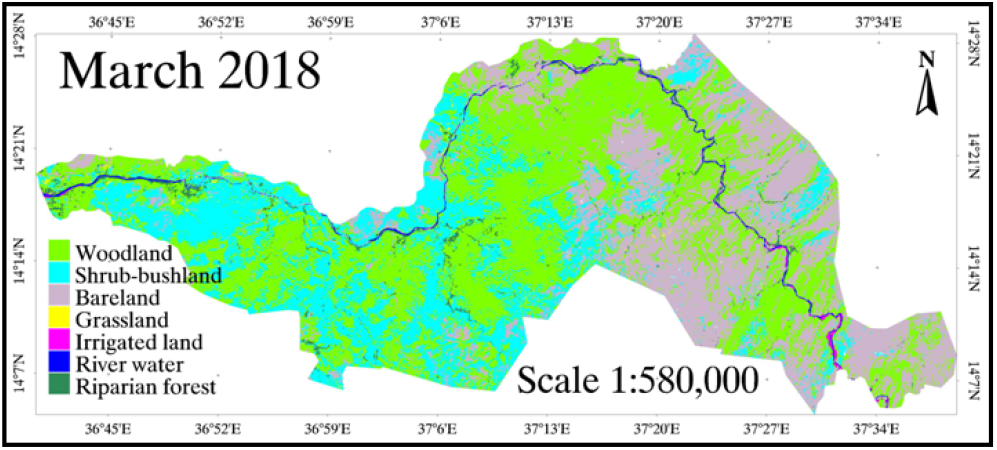
Dry season Land use land cover classes map derived from the Landsat imageries 2018

### Land use land cover change during the period 1988-2018

The magnitudes of land use land cover types, rate of change, and the annual rate of changes per year from 1988 to 2018 are summarized in Tables 7 and 8. Woodland area coverage during 1988 was the largest which accounts for 57.98% (1,251.88 km^2^) of the total area of the park. However, continuous declined of the woodland cover was observed in the three consecutive decades of the study period. But, the highest decline was observed during the second period (1998-2008) by 11.0% (124.8 km^2^) while the lowest decline was 9.57% (119.81 km^2^) between 1988 and 1998. In the entire period between 1988 and 2018 woodland was decreased by 29.38% (367.85 km^2^) at an average rate of 12.3 km^2^ (0.98%) per year. The increased legal and illegal settlement around the park, agricultural expansion, and firewood collection inside the park had a great impact on the destruction of woodland cover.

**Table 7:** Area extent of land use land cover types in 1988, 1998, 2008, and 2018

| Land use<br>land cover classes | 1988 |  | 1998 |  | 2008 |  | 2018 |  |
| --- | --- | --- | --- | --- | --- | --- | --- | --- |
|  | km<sup>2</sup> | % | km<sup>2</sup> | % | km<sup>2</sup> | % | km<sup>2</sup> | % |
| Woodland | 1,251.88 | 57.98 | 1,132.08 | 52.44 | 1,007.32 | 46.66 | 884.03 | 40.95 |
| Shrub-bushland | 374.91 | 17.36 | 402.36 | 18.64 | 436.82 | 20.23 | 507.18 | 23.49 |
| Riparian forest | 83.77 | 3.88 | 60.92 | 2.82 | 52.48 | 2.41 | 44.31 | 2.05 |
| Grassland | 371.03 | 17.18 | 417.52 | 19.34 | 496.35 | 22.99 | 532.34 | 24.66 |
| Agricultural land<sup>a</sup> | 0.00 | 0.00 | 64.79 | 3.00 | 95.57 | 4.45 | 118.35 | 5.48 |
| Water body | 48.30 | 2.24 | 49.22 | 2.28 | 34.59 | 1.60 | 35.02 | 1.62 |
| Bareland | 29.13 | 1.35 | 32.09 | 1.48 | 35.85 | 1.66 | 37.75 | 1.75 |
| **Total** | 2,159 | 100 | 2,159 | 100 | 2159 | 100 | 2159 | 100 |

<sup>a</sup>Crop cultivation area was absent in 1988 but agriculture in the park clearly started since 1993

**Table 8:** Magnitude and annual rate of change in different LULC categories of KSNP from 1988-1998, 1998-2008, 2008-2018 and 1988-2018*

| Land use<br>land cover<br>classes | 1988-1998 |  | 1998-2008 |  | 2008-2018 |  | Change<br>between<br>1988-2018 |  | Annual rate of<br>change/year<br>1988-2018 |  |
| --- | --- | --- | --- | --- | --- | --- | --- | --- | --- | --- |
|  | km<sup>2</sup> | % | km<sup>2</sup> | % | km<sup>2</sup> | % | (km<sup>2</sup>) | % | (km<sup>2</sup>) | (%) |
| Woodland | -119.81 | -9.57 | -124.8 | -11 | -123.3 | -12.2 | -367.85 | -29.38 | -12.3 | -0.98 |
| Shrub-bushland | 27.45 | 7.32 | 34.46 | 8.56 | 70.36 | 16.1 | 132.27 | 35.28 | 4.41 | 1.17 |
| Riparian forest | -22.85 | -27.3 | -8.78 | -14.4 | -7.83 | -15 | -39.46 | -47.11 | -1.32 | -1.57 |
| Grassland | 46.49 | 12.53 | 78.82 | 18.88 | 35.99 | 7.25 | 161.31 | 43.47 | 5.38 | 1.45 |
| Agricultural land | 64.80 | - | 30.78 | 47.5 | 22.78 | 23.8 | 118.36 | - | 3.94 | - |
| Water body | 0.92 | 1.89 | -14.63 | -29.7 | 0.42 | 1.23 | -13.29 | -27.51 | -0.44 | -0.92 |
| Bareland | 2.94 | 10.08 | 3.78 | 11.8 | 1.3 | 3.61 | 8.02 | 27.52 | 0.27 | 0.92 |

\*Negative sign (-) indicates the land use land cover class was decreasing in the entire time span

Shrub-bushland had the lowest coverage in 1988 accounting for 374.91 km^2^ (17.36%) and the highest during 2018 at 507.2 km^2^ (23.49%). The increase of shrub-bushland was maximum, 16.1% (70.36 km^2^) from 2008 to 2018. From 1988-2018 the shrub-bushland cover was increased by 132.27 km^2^ (35.28%) at a rate of 4.41 km^2^ (1.17%) per year due to illegal fire destructed the woodland cover for rainfed crop cultivation & continuous mass livestock grazing.

The riparian forest was declined consecutively during 1988-1998 and 1998-2008 by 27.3 km^2^ (22.85%) and 14.4 km^2^ (8.78%) respectively. However, in the 3^rd^ period (2008-2018) the forest slightly declined by 7.83 km^2^ (15%). In the three decades study, this class decreased by 39.46 km^2^ (47.11%) at an annual rate of 1.32 km^2^ (1.57%) per year. The depletion of Tekeze riverside riparian forest was due to extensive irrigated crops cultivation undertaken since 1993.

Grassland was computed the lowest coverage, 371.03km^2^ (17.18%) in 1988 unlike in 2018 with a maximum value, 532.34 km^2^ (24.66%). The highest expansion of grassland in the study area was 18.88% (79.82 km^2^) between 1998 and 2008. However, the increment was very small during 2008 and 2018, which counts 7.25% (35.99 km^2^). During the 1988-2018 period grassland (grazing land) expansion was showed by 43.47% (161.31 km^2^) at an annual rate of 5.38 km^2^ (1.45%) per year. Because of conversion of woodland to cultivation and destruction of woodland by firewood collection and charcoal production favored to grow seasonal grass and herbs.

Agricultural land (irrigated and rainfed crops) clearly observed a highly expanded as compared to the other land cover classes. Crop cultivation inside the park was absent in 1988, however, it was started after 1993 and continued as for 1998, 2008, and 2018 which has extent cover about 3%, 4.43%, and 5.48% respectively. The highest expansion of cultivation was observed between 1993 and 1998 by 64.8 km^2^ following 47.5% (30.78 km^2^) from 1998 to 2008. However, the expansion was relatively small, 23.8% (22.78 km^2^) between 2008 and 2018. During 1993-2018 cultivation was increased by 118.36 km^2^ at an annual rate of 3.94 km^2^ per year. This expansion of agriculture was due to the expansion of the settlement program surrounding the park and extensive rain-fed crop cultivation by illegal highland settlers of the region.

Water bodies’ coverage in the years 1988, 1998, 2008, and 2018 was 2.24%, 2.28%, 1.6%, and 1.62% respectively from the total area of the park. In three decades (1988-2018) water negatively changed by 13.29 km^2^ (27.51%) with the annual rate of 0.44 km^2^ (0.92%) per year.

Bareland’s highest expansion of change rate was 11.8% (3.78 km^2^), during 1998 and 2008 followed by 10.08% (2.94 km^2^) and 3.61% (1.3 km^2^) from 1988 to 1998 and 2008 to 2018 respectively. In the whole study period, bare land expansion was 27.52% (8.02 km^2^) with the annual rate of increase by 0.27 km^2^ (0.92%) per year between 1988 and 2018. However, based on the seasonal comparison of March 2018 and October 2018, bareland cover tremendously expanded from wet months to drier months (Fig 6).

**Fig 6:**
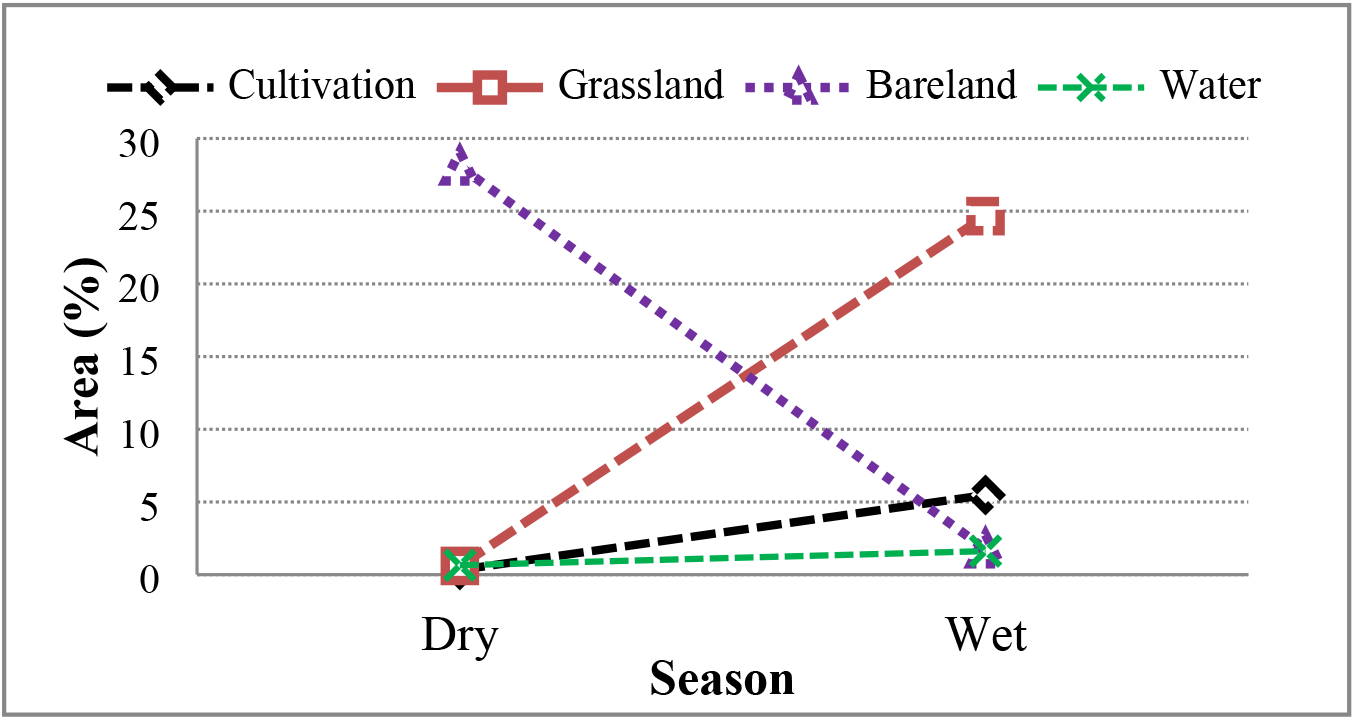
Seasonal landuse landcover classes (Bareland, cultivation, grassland, and water) cover change of Kafta-sheraro national park in March and October 2018

In addition to temporal variation of LULC change, some classes varied their proportion and rate of change seasonally (dry and wet). Thus, the comparison between the wet season (October 2018) and dry season (March 2018) satellite imageries of LULC change classification of KSNP was computed (Fig 4 and 5). The result indicated that the land cover class of bareland increased by 26.3% (568.46 km^2^) from the wet season towards the drier season because the total rain-fed crop field and grazing areas were changed into bare ground during long dry months. In contrast, cultivation (rainfed crops), grazing land, and water decreased by 5.15% (111.26 km^2^), 24.06% (519.5 km^2^), and 0.97% (21 km^2^) respectively from wet months to the direction of drier months (Fig 6). The pronounced change in bareland occurred due to the harvesting of rainfed crops and dried and removed pasture during the prolonged dry months.

### Temporal and seasonal change of NDVI

The area of the study increasingly exercised illegal fire, extensive agriculture, charcoal production, and other related human-induced drivers of LULC change. These factors change and negatively affected the vegetation resources of the park. To detect vegetation cover change of thirty years period, normalized difference vegetation analysis of 1988 and 2018 wet and dry seasons’ satellite sensors were utilized. Fig 7 indicates the NDVI thresh hold based difference of dry (mid of March) and wet (mid of October) months of the year 1988 and 2018. The pixels count of the wet season showed a higher NDVI value of 0.85 in 1988 and 0.84 in 2018 from June to mid-October; the park areas are dominated by vegetated woodland, shrubland, and riparian (Riversides) vegetation. The vegetation occurrence is relative, denser in the Western, Northwest, and Central parts of the park while areas of small portion are found in Eastern and Southern parts. During the dry season, from the end of December to the end of March, the maximum NDVI value was 0.64 in 1988 and 0.49 in 2018. Similarly, the average NDVI value in dry 2018 decreased to 0.047+0.064 (mean+SD) whereas in wet season reduced to 0.066+0.096. During the dry season dominant vegetated areas concentrated on the sides of water points (hereafter Tekeze Riversides and its tributary rivers) and the Eastern riversides irrigated fruit plantations of the park. Areas of low NDVI value were observed in the dry season because during the wet season a section of the park area was used as rainfed crop cultivation, grazing, and areas deforested by illegal fire and these situations clearly observed while we conducted the field training.

**Fig 7:**
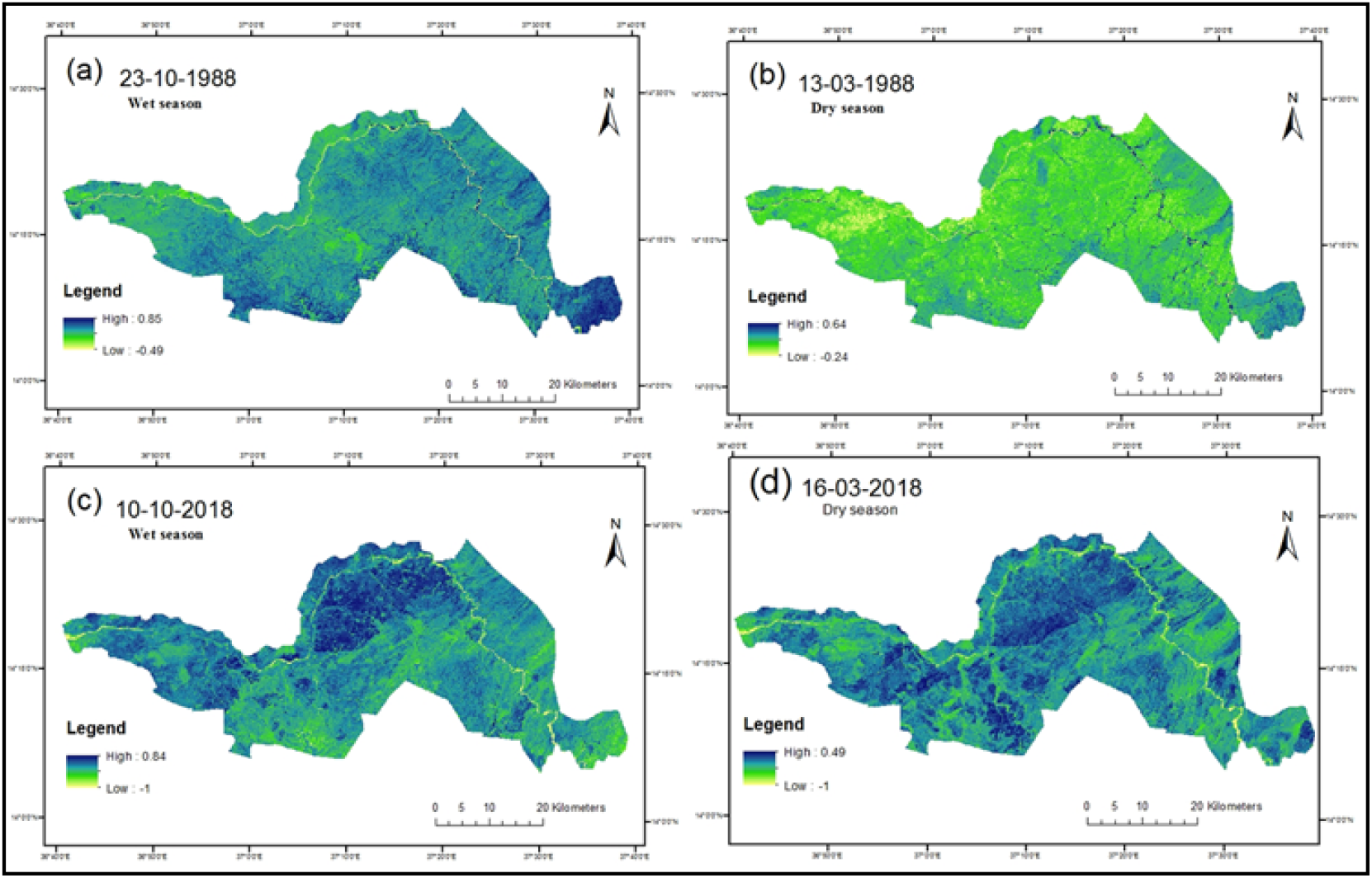
Seasonal varibility of NDVI (greeness status) in Kafta-sheraro national park (KSNP) for wet season (**a** and **c**) and dry season (**b** and **d**) during the year 1988 and 2018 satellite images

The present reclassify analysis of NDVI also showed a significant change in areas of vegetation cover (Fig 8 and Table 9). The high-moderate density vegetation cover in 1988 was about 66.67%. However, the magnitude of its cover in 2018 was declined to 45.2%. In the entire period of 1988-2018 high-moderate density vegetation reduced by 464.6 km^2^. The sparse vegetation covered 29.8% from the total area of the park in 1988 but expanded to 49.6% in 2018. Sparse vegetation coverage was increased by 428.1 km^2^ from the total area of the park between 1988 and 2018. Moreover, the coverage of non-vegetation was increased from 3.5% in 1988 to 5.2% in 2018. Non-vegetation showed an expansion of 36.5 km^2^ from 1988 to 2018 (Table 9).

**Fig 8:**
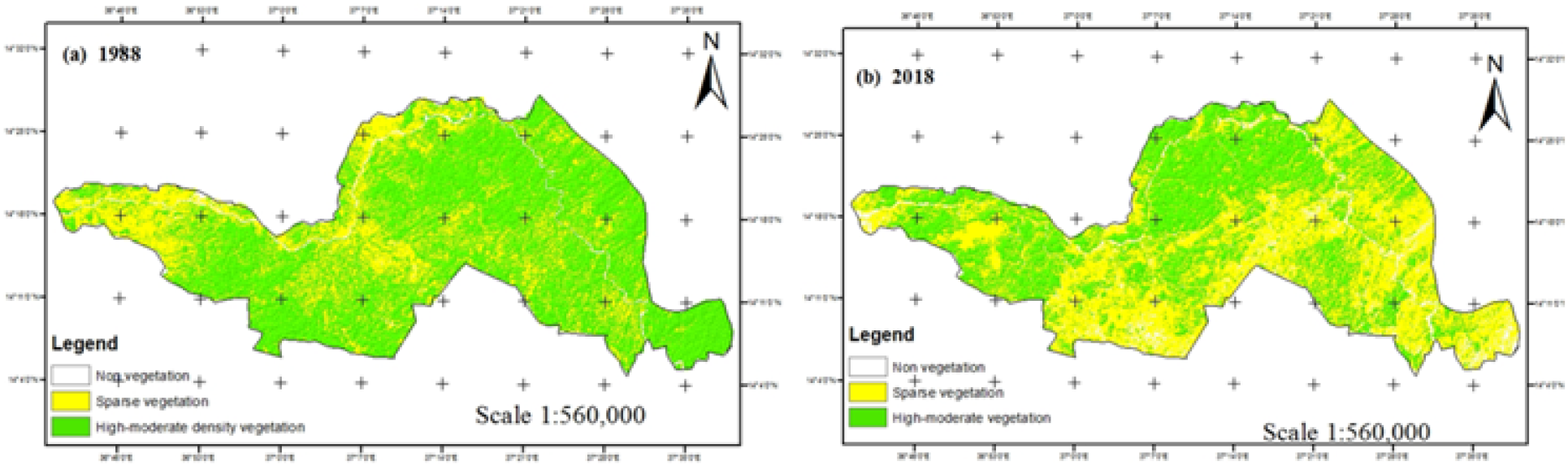
Vegetation covers change map of Kafta-Sheraro National Park in 1988 (a) and 2018(b)

**Table 9:** Normalized difference vegetation index (NDVI) of land cover change (area in km^2^ and %) in Kafta-sheraro national park between 1988 and 2018

| Land cover class | 1988 |  | 2018 |  | 2018-1988 |
| --- | --- | --- | --- | --- | --- |
|  | km<sup>2</sup> | % | km<sup>2</sup> | % | Change (km<sup>2</sup>) |
| High-moderate density vegetation | 1439.4 | 66.67 | 974.8 | 45.2 | -464.6 |
| Sparse vegetation | 643.5 | 29.8 | 1071.6 | 49.6 | +428.1 |
| Non-vegetation | 76.115 | 3.5 | 112.6 | 5.2 | +36.5 |
| Total | 2,159 | 100 | 2,159 | 100 | - |

### Seasonal NDVI -precipitation/temperature relationship

According to the wet and dry season’s Landsat data, the inter-annual variation of NDVI during the 10 years period (2007-2016) was computed (Fig 9a). The mean NDVI of dry and wet seasons showed variable trends and differ significantly at (p<0.05) over ten years period. Relative to the wet season, the dry season NDVI indicated a decreasing trend. The minimum and maximum wet season mean NDVI values were 0.21 in 2012 and 0.33 in 2007 respectively and intermediate values appeared in the rest of the years. In the dry season, the maximum mean value NDVI was a score for 2007, 2008, and 2013 while minimum value occurred in 2016. In contrast, the rainfall and temperature did not show significant variation in the specified period (Fig 9 b-c). Therefore, the variation and slight decreasing trends in NDVI might occur due to several activities of the local communities like extensive woodland conversion to cultivation and extinguishing of illegal fire, and the dominant nature of scattered woodland vegetation composition of the park.

**Fig 9:**
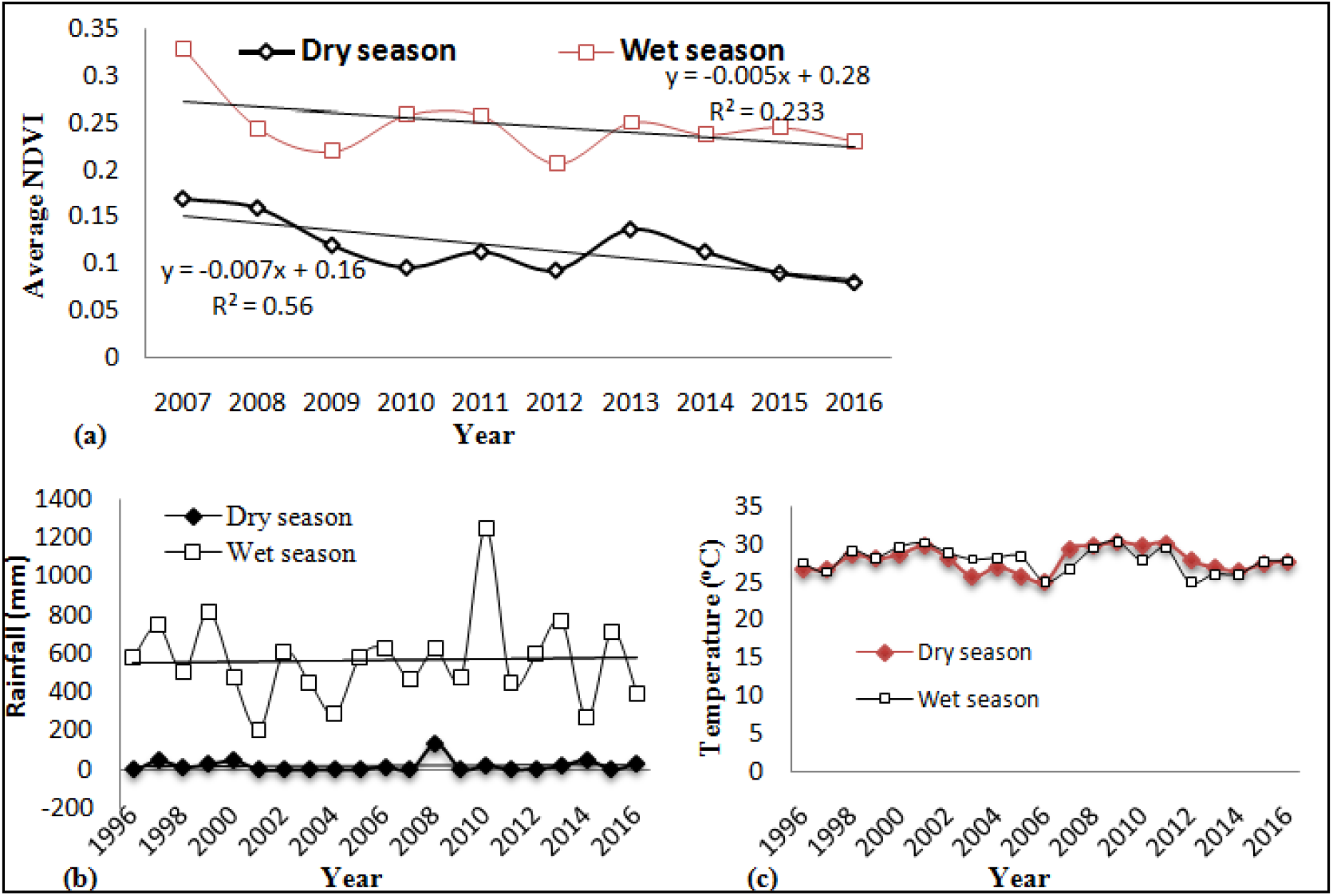
Dry and wet season inter annual variation in a seasonal mean NDVI between 2007 and 2016(**a**), seasonal annual rainfall (**b**), and seasonal mean temperature (**c**) in KSNP

The relationship between NDVI and climate variables (rainfall and temperature) was analyzed in KSNP for ten years period (2007 to 2016) because satellite image data was absent and some missing values of daily rainfall and temperature before 2007. Fig10a-d summarizes the statistical analysis between seasonal mean NDVI, mean annual rainfall (values scale in Log10 to fit NDVI values), and mean temperature between 2007 and 2016. Even if there is a change in seasonal rainfall and temperature, the correlation relationship of wet and dry seasons NDVI with these two climate variables did not show statistically significant (p>0.05).

**Fig10:**
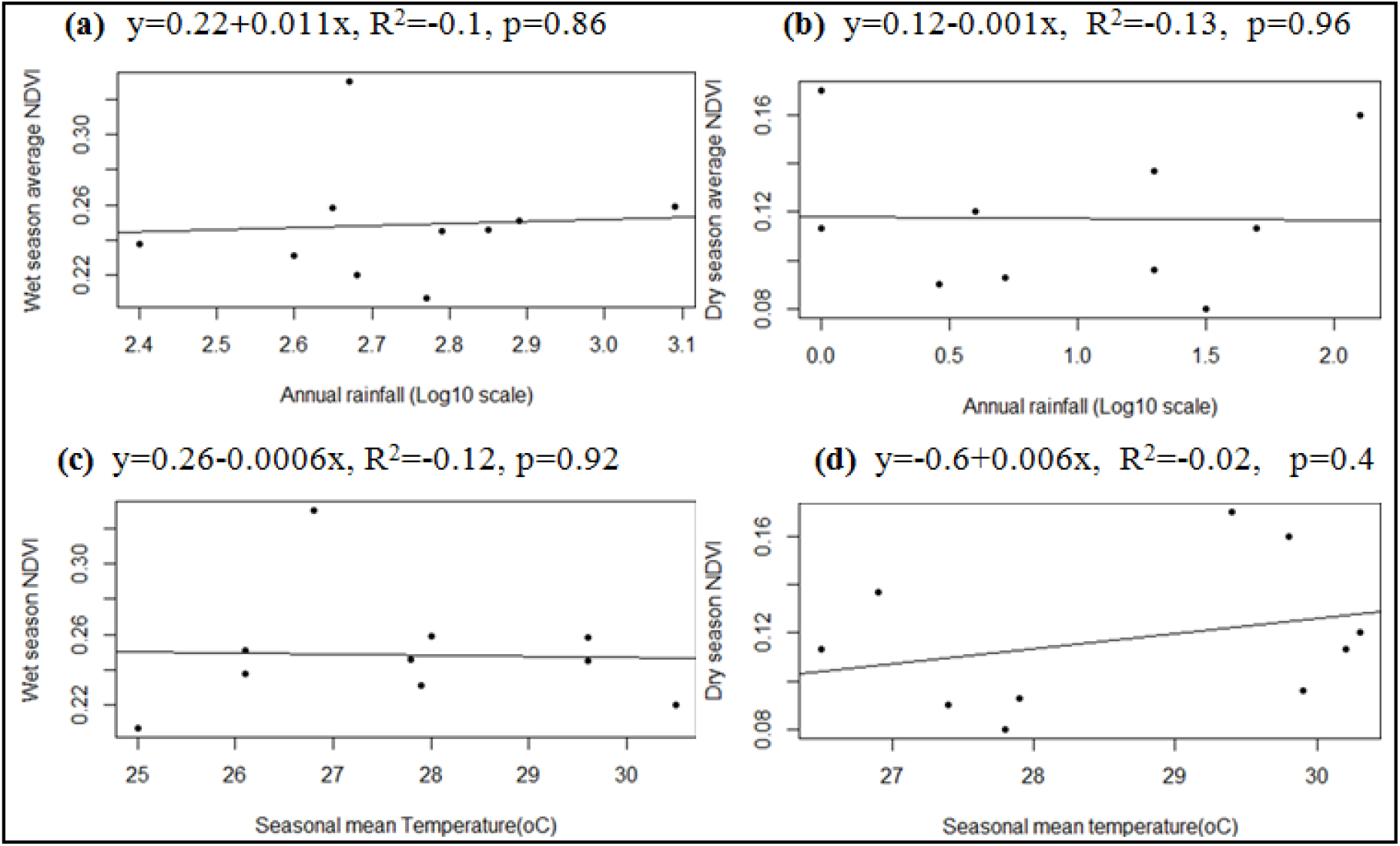
Regression analysis of seasonal mean NDVI trend and relationship with (a and b) wet and dry season’s rainfall (mm) and temperature (°C) (c and d) between 2007 and 2016

### Local community perception on drivers of LULC change

#### Demographic and socio-economic information of sampled households

**Table 10** provides a summary of the demographic and socio-economic features of the sampled households. The age of the respondents ranged from 22 to 75 years, with an average of 44.7 years, and above fifty percent of respondents’ age category (22-41 years) is under the productive region of human resources. About 71.7% of the interviewees came from other areas and resettled after 1991 and the rest existed before in the area. About 74% of the sampled households were male. The household size ranged from two persons to 8 persons, with an average of ~5 persons. The farm size of the respondents varied from 1 to 10 hectares, with an average of 3.9 hectares. With respect to their education status, 74.2% of the respondents were attended formal and 25.8% had attended informal education.

**Table 10:** Socio-economic general characterstics of sampled household (N=395) in the study

| House hold characteristics | Catagories | Value in | Estimated value |
| --- | --- | --- | --- |
| Gender | Male | % | 74.0 |
| Age | - | Years | min:22; mean: 44.7 ; max:75 |
| Ethnic category | Tigraway | % | 86.0 |
|  | Kunama | % | 14.0 |
| Family size | 2-5 ;6-8 | Number | min: 2; mean: 5; max:8 |
| Education status | Formal (1-10<sup>th</sup> grade) | % | 74.2 |
| Resettlement status (1991-2003) | - | % | 71.7 |
| Distance of settlement from park | 5-10; 11-15; >15 km | km | min: 5; mean: 12.6 ; max:21 |
| Respondents alternative income | - | % | 12.9 |
| Energy for cooking | Firewood | % | 94.7 |
| Farmland holding size | 1-3; 4-7; >7 ha | hectare | min:1; mean: 3.9; max:10 |
| Crop use (consumption and sale) | - | % | 74.0 |
| Source of livelihood & income | Crop production | rank | 1 |
| Land tenure (land use permits) | Legal | % | 86.0 |

Around three fourth of the sampled households were engaged in farming of crop production and mixed crop and livestock. However, a small portion of the respondents was involved only in livestock rearing and other additional activities couple with farming. Crop production was ranked as the topmost important source of livelihood/ income and followed by mixed crop and livestock farming in the district (**Table 11**). The most important types of crop produced in the study area were both rainfed, such as *Sesamum indicum, Sorghum bicolor, Eleusine coracana, Eragrostis tef*, and *Zea mays* L. whereas irrigated crops, like *Muza species, Mangifera indica, Carica papaya, Allium cepa, Allium sativum, Solanum tuberosum*, and *Capsicum annuum*.

**Table 11:** The main livelihood and economic activities ranked by respondents (N=395^$^)

| Activities | No of respondents | Percentage |
| --- | --- | --- |
| Crop production (rainfed and irrigated) | 197 | 50.0 |
| Livestock rearing | 25 | 6.3 |
| Mixed crop and livestock | 173 | 43.7 |
| Private and government employee | 22 | 5.5 |
| Self business | 17 | 4.3 |
| Wood product (charcoal & fire wood collection) | 5 | 1.3 |
| Gold mining | 7 | 1.8 |

<sup>§</sup> The total number of respondents was 395, however, over counts are predictable due to multiple response of households to questions

#### Local community response on trends of LULC variables

Local community perception on major LULC change types and other associated variables of change showed statistical significance (p<0.001). The participants were efficiently aware that woodland and riparian forest (Tekeze river side vegetations of the park) significantly declined in the whole studied period (p<0.001). The local communities had given confidential evidence for the decline of about 90.3% woodland and 88.7% riparian vegetation of KSNP and its surrounding. In contrast, around 57.3 % of the respondents recognized that distance from water source to settlement had shown constant trends. However, agricultural land, wet season grazing area (grassland), bare land, resettlement, and road access to the park had significantly increased throughout the studied periods (p<0.001) (**Fig 11**).

**Fig 11:**
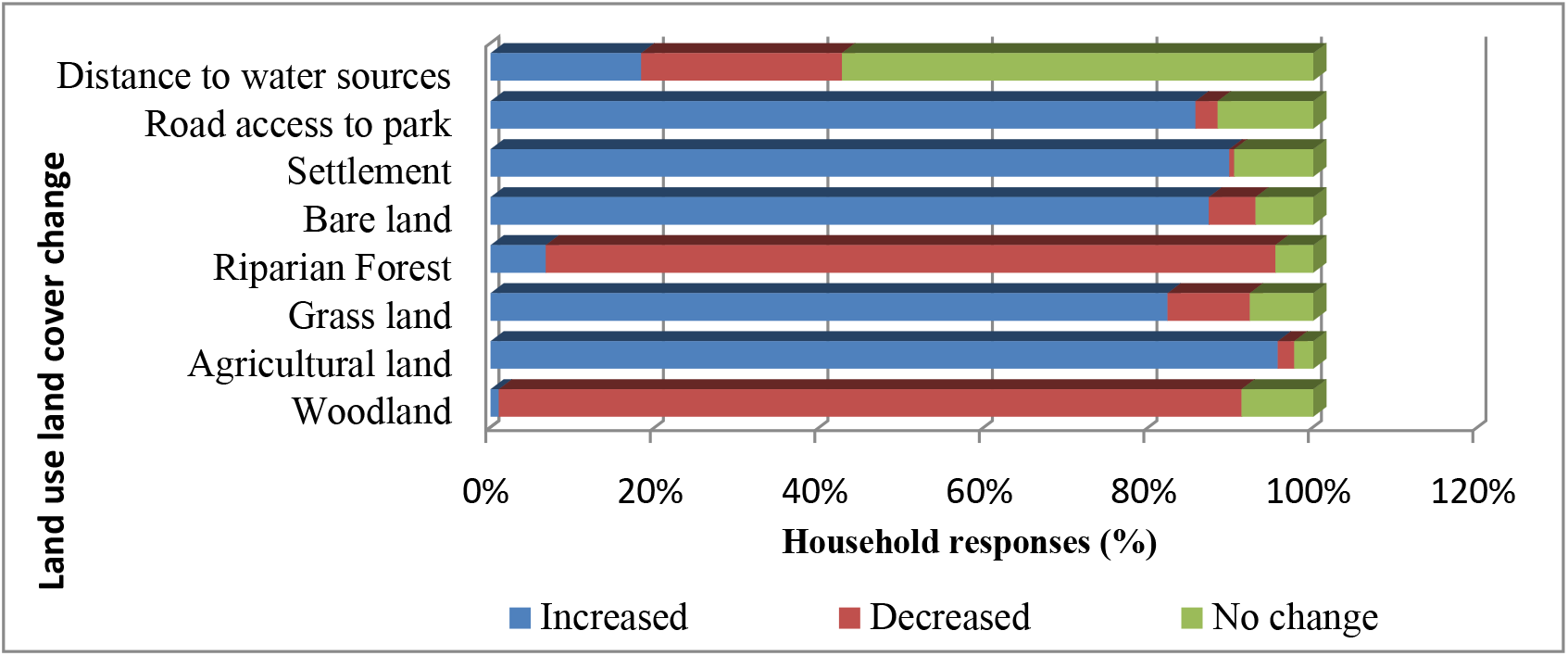
Respondents awarness towards the trend of land use land cover variables (N=395)

**Fig 12:**
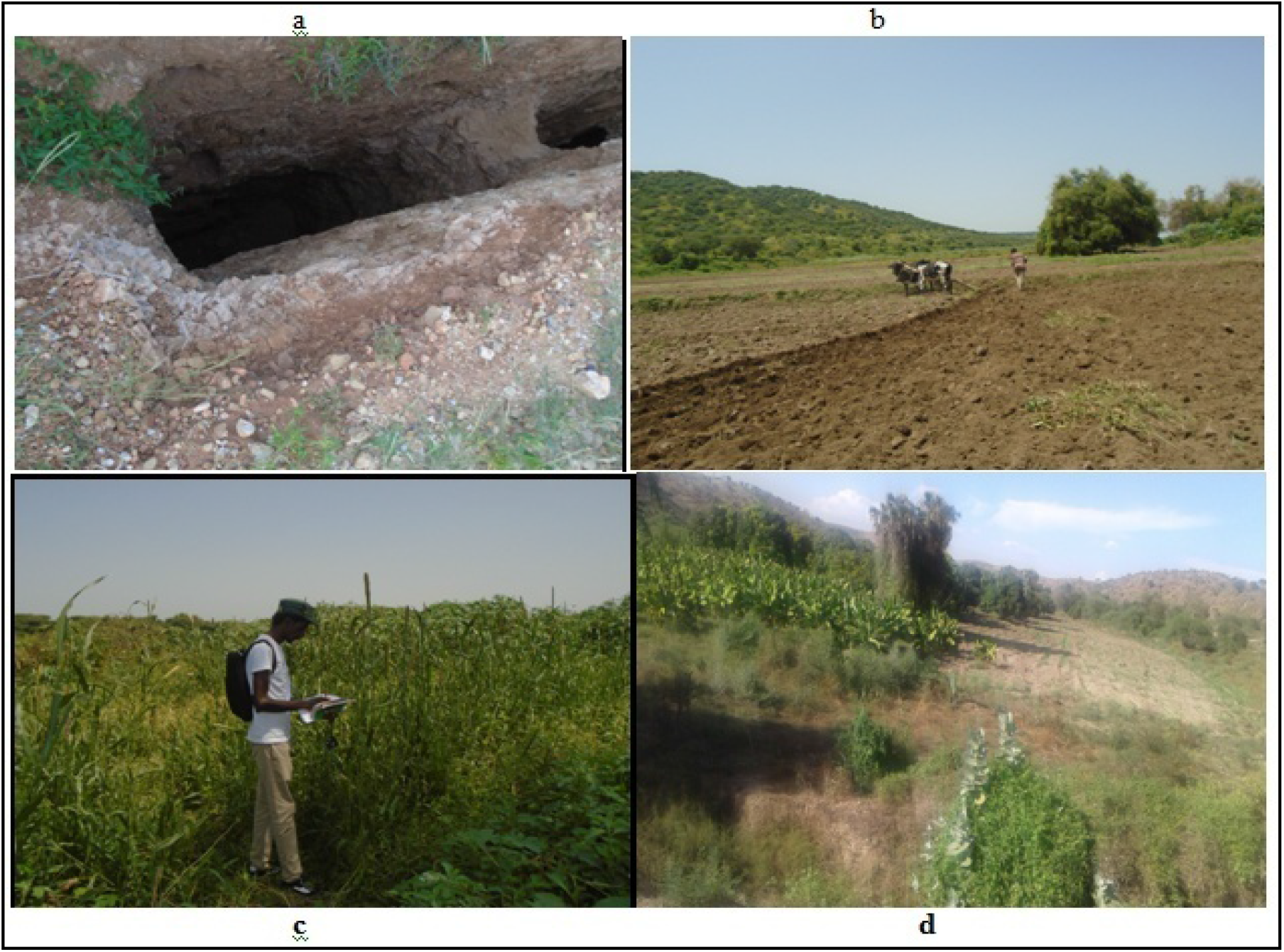
Traditional gold mining activities (a) preparing land for planting (b) and cultivations of Banana and cereal crops (c and d) along Tekeze riversides of Kafta-Sheraro National Park (Photo by Fitsum Temesgen, 2018-2019)

#### Main (immediate) drivers of LULC changes in KSNP

During the surveyed period the participants identified 13 pronounced factors as key drivers of the observed LULC change in KSNP. Increasing the expansion of legal and illegal settlements, cultivated land, illegal fire following encroachment by cultivation, seasonal grazing, firewood collection, and traditional gold mining were prioritized the top significantly (p<0.001) ranked and immediate drivers of LULC in the park (**Table 12**). Likewise, from key informant interviews and focus group discussions (FGDs), similar feedbacks were also exhibited and they were strengthened that; expansion of settlements and agriculture, firewood collection, charcoal production, and land tenure (administration) problems were identified as the main causes of LULC change in the study area.

**Table 12:**
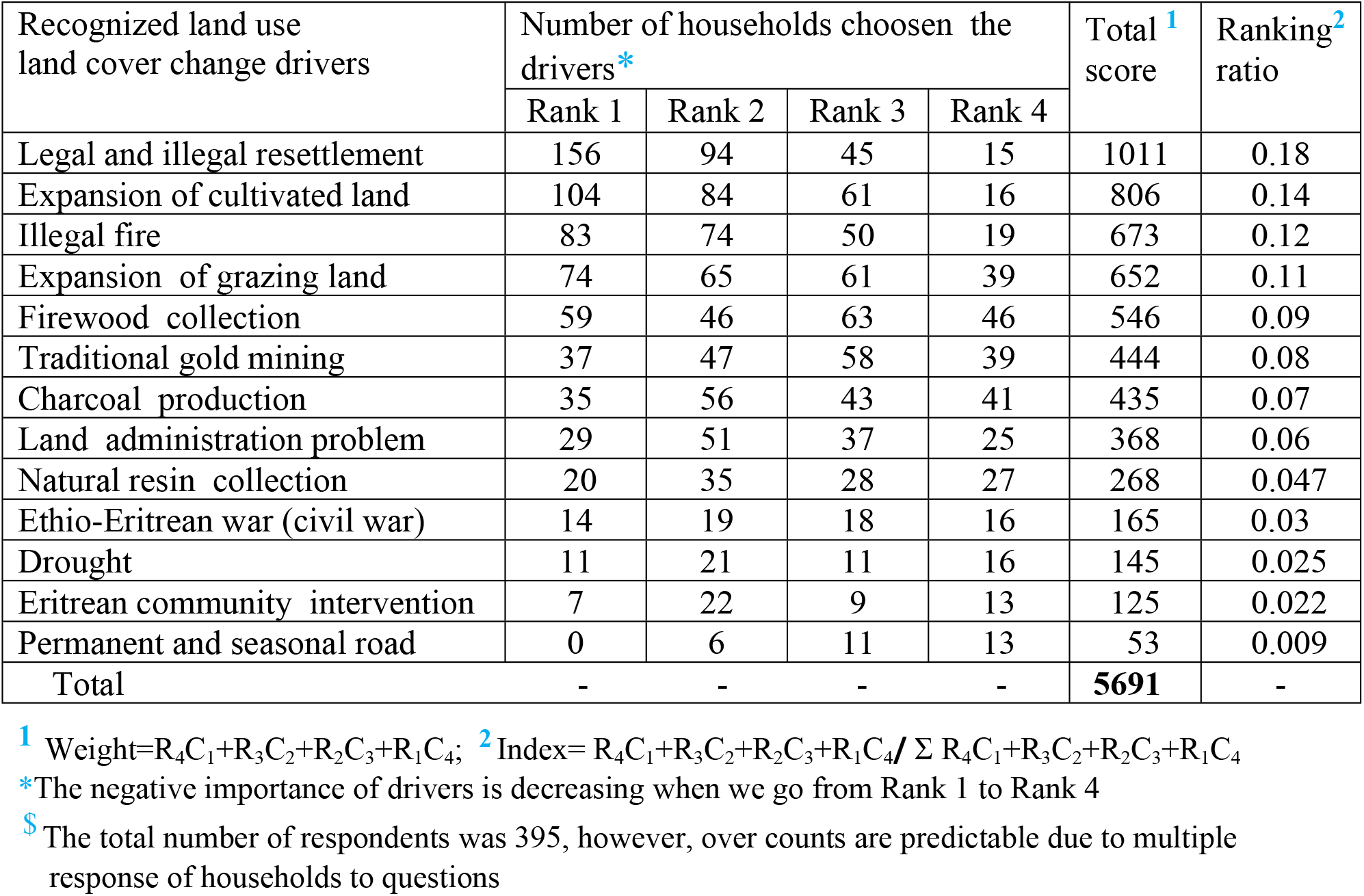
Main drivers of LULC change recognized by local communities’ perception in Kafta-sheraro National park (N=395^$^)

#### Socio-economic characterstics vs respondents’ awareness on drivers

To test the statical significance, logistic regression analysis checked (p<0.05) on the top five recognized drivers namely: resettlement, expansion of agriculture, fire, expansion of grazing land, and traditional gold mining at the household level (**Table 13**). Seven demographic and socio-economic explanatory (independent) variables were utilized for the whole analysis. The result of the analysis indicated that settlement duration (p=0.01) and distance from settlement to park (p=0.04) significantly and negatively influenced the sampled households’degree of awareness on expansion of agriculture and grazing land. Additionally, respondents’ high awareness of expansion of cultivated land, settlement, illegal fire, expansion of grazing, and gold mining was influenced significantly (p=0.001) and negatively by education level of the interviewees. The perception of respondents on expansion of grazing land and expansion of settlement (resettlement) also negatively influenced by individual household size and agricultural land size occupied (p=0.045). The rest explanatory variables such as age category, gender, number of families, and the landholding size of respondents were not significantly influenced the perception of respondents on the five below listed drivers of LULC change (**Table 13**).

**Table 13:** The household demographic and socio-economic status impacts on their attitude towards the top five recognized drivers of LULC change in KSNP

| **1. Expansion of cultivated land (driver)** |  |  |  |  |  |  |  |  |
| --- | --- | --- | --- | --- | --- | --- | --- | --- |
| *Independent variables* | B | S.E | Wald | df | p-value | Exp(B) | 95% C.I.for EXP(B) |  |
|  |  |  |  |  |  |  | Lower | Upper |
| Age | 0.62 | 0.45 | 1.87 | 1 | 0.17 | 1.86 | 0.76 | 4.55 |
| Gender (1=male) | 0.20 | 0.31 | 0.42 | 1 | 0.52 | 1.22 | 0.66 | 2.24 |
| Education level (1=Informal education )* | -4.78 | 1.08 | 19.43 | 1 | 0.00 | 0.008 | 0.001 | 0.07 |
| Education level (2=Formal:1-8 & 9-10 grade) | 0.05 | 0.41 | 0.02 | 1 | 0.89 | 1.06 | 0.47 | 2.37 |
| House hold size | -0.73 | 0.41 | 3.08 | 1 | 0.08 | 0.48 | 0.21 | 1.09 |
| Settlement duration* | -0.79 | 0.31 | 6.60 | 1 | 0.01 | 0.45 | 0.25 | 0.83 |
| Agricultural land size | -0.41 | 0.56 | 0.54 | 1 | 0.46 | 0.66 | 0.22 | 1.97 |
| Distance from settlement to park (1=5-10 km) | -0.22 | 0.29 | 0.58 | 1 | 0.44 | 0.80 | 0.45 | 1.41 |
| **Table 13. Continued** |  |  |  |  |  |  |  |  |
| **2. Resettlement (driver)** |  |  |  |  |  |  |  |  |
| *Independent variables* | B | S.E | Wald | df | p-value | Exp(B) | 95% C.I.for EXP(B) |  |
|  |  |  |  |  |  |  | Lower | Upper |
| Age | -0.44 | 0.42 | 1.10 | 1 | 0.29 | 0.64 | 0.28 | 1.46 |
| Gender (1=male) | -0.47 | 0.29 | 2.70 | 1 | 0.10 | 0.62 | 0.35 | 1.09 |
| Education level (1=Informal education )* | -2.10 | 0.54 | 15.21 | 1 | 0.00 | 0.12 | 0.04 | 0.35 |
| Education level (2=Formal:1-8 & 9-10 grade) | -0.09 | 0.41 | 0.05 | 1 | 0.82 | 0.91 | 0.41 | 2.04 |
| House hold size | 0.55 | 0.38 | 2.12 | 1 | 0.14 | 1.73 | 0.83 | 3.64 |

| Settlement duration | 0.22 | 0.29 | 0.60 | 1 | 0.44 | 1.25 | 0.71 | 2.19 |
| --- | --- | --- | --- | --- | --- | --- | --- | --- |
| Agricultural land size* | -1.23 | 0.66 | 3.46 | 1 | 0.045 | 0.29 | 0.08 | 1.07 |
| Distance from settlement to park (1=5-10 km) | 0.18 | 0.27 | 0.43 | 1 | 0.51 | 1.19 | 0.70 | 2.04 |
| **3. Human induced fire (driver)** |  |  |  |  |  |  |  |  |
| *Independent variables* | B | S.E | Wald | df | p-value | Exp(B) | 95% C.I.for EXP(B) |  |
|  |  |  |  |  |  |  | Lower | Upper |
| Age | 0.25 | 0.44 | 0.34 | 1 | 0.56 | 1.29 | 0.55 | 3.03 |
| Gender (1=male) | 0.24 | 0.30 | 0.64 | 1 | 0.42 | 1.23 | 0.70 | 2.30 |
| Education level (1=Informal education )* | -3.31 | 0.64 | 26.74 | 1 | 0.00 | 0.04 | 0.01 | 0.13 |
| Education level (2=Formal:1-8 & 9-10 grade) | 0.16 | 0.41 | 0.15 | 1 | 0.70 | 1.17 | 0.52 | 2.63 |
| House hold size | -0.05 | 0.39 | 0.02 | 1 | 0.89 | 0.95 | 0.44 | 2.03 |
| Settlement duration | 0.17 | 0.30 | 0.33 | 1 | 0.56 | 1.19 | 0.66 | 2.15 |
| Agricultural land size | -0.29 | 0.54 | 0.28 | 1 | 0.59 | 0.75 | 0.25 | 2.18 |
| Distance from settlement to park (1=5-10 km) | -0.22 | 0.25 | 0.61 | 1 | 0.43 | 0.80 | 0.46 | 1.40 |
| **4. Expansion of grazing (driver)** |  |  |  |  |  |  |  |  |
| *Independent variables* | B | S.E | Wald | df | p-value | Exp(B) | 95% C.I.for EXP(B) |  |
|  |  |  |  |  |  |  | Lower | Upper |
| Age | 0.57 | 0.45 | 1.61 | 1 | 0.20 | 1.77 | 0.73 | 4.31 |
| Gender (1=male) | 0.03 | 0.30 | 0.01 | 1 | 0.93 | 1.03 | 0.56 | 1.86 |
| Education level (1=Informal education )* | -3.83 | 0.82 | 21.97 | 1 | 0.00 | 0.02 | 0.004 | 0.11 |
| Education level (2=Formal:1-8 & 9-10 grade) | 0.31 | 0.41 | 0.59 | 1 | 0.44 | 1.37 | 0.61 | 3.03 |
| House hold size * | -0.77 | 0.42 | 3.45 | 1 | 0.045 | 0.46 | 0.20 | 1.04 |
| Settlement duration | -0.02 | 0.31 | 0.01 | 1 | 0.93 | 0.97 | 0.53 | 1.77 |
| Agricultural land size | 0.23 | 0.58 | 0.16 | 1 | 0.69 | 1.26 | 0.40 | 3.92 |
| Distance from settlement to park(1=5-10 km)* | -0.56 | 0.29 | 3.78 | 1 | 0.04 | 0.57 | 0.32 | 1.00 |
| **5. Traditional gold mining (driver)** |  |  |  |  |  |  |  |  |
| *Independent variables* | B | St | Wald | df | p-value | Exp(B) | 95% C.I.for EXP(B) |  |
|  |  |  |  |  |  |  | Lower | Upper |
| Age | -0.07 | 0.38 | 0.05 | 1 | 0.85 | 0.93 | 0.44 | 1.96 |
| Gender (1=male) | 0.08 | 0.25 | 0.09 | 1 | 0.76 | 1.08 | 0.65 | 1.78 |
| Education level (1=Informal education )* | -1.20 | 0.47 | 6.54 | 1 | 0.01 | 0.30 | 0.12 | 0.75 |
| Education level (2=Formal:1-8 & 9-10 grade) | -0.72 | 0.43 | 2.80 | 1 | 0.09 | 0.48 | 0.21 | 1.13 |
| House hold size | 0.06 | 0.34 | 0.03 | 1 | 0.86 | 1.06 | 0.54 | 2.06 |
| Settlement duration | -0.05 | 0.26 | 0.04 | 1 | 0.83 | 0.95 | 0.57 | 1.58 |
| Agricultural land size | 0.80 | 0.52 | 2.31 | 1 | 0.13 | 2.22 | 0.79 | 6.24 |
| Distance from settlement to park (1=5-10 km) | -0.22 | 0.25 | 0.76 | 1 | 0.38 | 0.80 | 0.49 | 1.31 |

\*=significant at 5% (0.05), S.E=standard error, C.I=confidence interval)

## Discussion

### Seasonal variation of accuracy assessment

The wet and dry season’s classification accuracy assessment of Kafta Sheraro National Park (KSNP) is considered reliable and acceptable agreement for the classified image of 2018. Wet season overall accuracy (86.9%) and Kappa Coefficient (0.845) is higher than Central Rift-Valley of Ethiopia [**99**], protected and communal areas of Namibia [**54**], Quirimbas National Park, Mozambique [**20**], and Little Zab River basin [**100**]. Similarly, the dry season of overall accuracy (91.96%) and Kappa coefficient (0.90) value of the study is also higher than Bale Mountain National Park [**42**], Maputaland-Pondoland-Albany Biodiversity hotspot [**30**], Tarangire and Katavi parks [**101**], in hilly areas of China [**102**], and semi-arid India [**46**].

Comparatively, the dry season accuracy in 2018 was quantified higher than the wet season and these results are in agreement with the LULC study in the tropical semi-arid areas [**103**]. These authors tested and approved on wet and dry seasonal variation of Landsat-8 image and their classification accuracy exhibited an increasing trend from wet to drier seasons. In our study low classification accuracy or higher confusion error of the wet season, grassland, and woodland cover in 2018 was mostly caused by confusion with other related land cover classes. For example, in the wet season grass is often confused with cultivation, and woodland is confused with shrub-bushland due to the limited spatial resolution and image quality. [**103**] also observed the wet season grassland had low classification accuracy. During the wet season, there is a high vegetative cover of crops and natural vegetations which create confusion or made difficult it to differentiate among them [**46**]. A study in Burkina Faso also revealed the highest classification accuracy in the dry season and lowest in the wet season [**104**].

### Trends of land use land cover change

The LULC change analysis from 1988-2018 in Kafta-sheraro national park resulted in a change in land use land cover patterns either positively or negatively, this most likely affects the whole habitats of the park. The park experienced extensive LULC change due to increased settlement following the expansion of farming activities. A rapid reduction of woodland and the riparian forest occurred. The highest decline of woodland occurred in the second period (1998-2008) whereas riparian forest was in the first period (1988-1998). Because these periods took place widespread expansion of dry season irrigated land and wet season rainfed crops. In contrast, the dry season of bareland, agricultural land, grassland, and shrub-bushland was highly expanded throughout the studied period (1988-2018). The result is similar to a study in Babile elephant sanctuary from 1977 to 2017, woodland and riparian forests decreased whereas agricultural land and bareland continuously increased [**43**]. Farmland expansion as the expense of forest and woodland decline was also reported [**105,106**]. More studies from the central Rift Valley of dryland revealed areas of cropland doubled, grass/grazing land and bareland increased as the cost of woodland destructed between 1986 and 2016 [**85,99,107**]. In advance to cultivated land, a remarkable increasing trend was showed mainly at the expense of forest cover [**108**]. According to [**109**] expansion of grassland, cultivation, and settlement as a cost of natural forest cover destruction was suggested. On the other hand, woodland by 34.6% and forest by 59.9% decreased while cultivation expanded by 15.16 km^2^/year [**110**]. In three decades (1957 to 2014) the forest area of Gelda district decreased while shrub, farmland, and bareland expanded [**87**]. In contrary to the present study grassland and bareland were declined in Gozamin district between 1986 and 2018 [**111**]. Therefore, including the current study (KSNP), in all the listed case studies in Ethiopia showed agricultural land has expanded intensively at the expense of dense forest and woodland cover decline.

The conversion of woody vegetation to cultivation was also broadly reported in Kenya [**112, 113**]. Moreover, a study in semi-arid Karamoja of Uganda and MPA of South Africa showed expansion of crop cultivation and heightened encroachment of shrub-bushland as the cost of woodland decline [**30,114**]. Likewise, [**115**] revealed that bareland areas extremely increased in Malawi as the expense of forest decline. Likewise, the LULC change studies in a forest of Kenya [**116**]; Cameron highlands [**117**]; semi-arid region of India [**46**]; Godavari basin of India [**118**]; Central Vietnam [**119**]; and in Islamabad of Pakistan [**120**] revealed a pronounced decrease of forest areas while an extensive expansion of agricultural land. A similar study in the Kalahari woodland savanna biome of Namibia showed expansion of agriculture at the expense of woodland Savannah reduction [**121**]. From 1979-2017, Quirimbas National Park has lost 41.67% of forest and changed to other human-made LULC [**20**]. In central Kenya farmland and bare land increased by 160.45% and 73.2% respectively as a cost of forest decline [**122**].

### Trends of temporal and seasonal NDVI

The LULC change analysis of Kafta-sheraro national park (KSNP) took for dry and wet seasons separately. In both seasons land cover classes of area coverage and in vegetating status differences were observed. Bareland was very small in the wet season but increased towards drier months whereas grassland (grazing land) and agriculture were relatively small in the dry season but increased in the wet season. LULC change variation occurred between the dry season irrigated crops and the wet season rainfed crops [**103**]. The authors suggested that during the dry season bareland is higher and higher while in the wet season is very small. The seasonal variation of average NDVI between wet and dry seasons was observed and reported in semi-arid areas [**123,124**]. Likewise, in semiarid areas of Uganda, the mean NDVI of the wet season was higher than the dry season [**114**]. In our study the result of NDVI also indicated there was a significant change in vegetation cover; the amount of high-moderate density vegetation cover was declined by 21.47% while sparse vegetation cover was increased by 19.8% from the total area of KSNP. On the other hand, non-vegetation cover increased by 36.5 km^2^ between 1988 and 2018. In agreement with the present study, the dense vegetation cover was declined by 26.1% whereas non-vegation increased by 14.3 km^2^ between the periods of 2000 and 2018 [**110**]. In Kafta-humera district (surrounding the park) woodland vegetation converted by cropland leads to expand bare ground during the dry season [**40**].

The NDVI indicated a pronounced decline both in wet and dry seasons of the study area with a little effect of rainfall and temperature. In the present study, the variation of seasonal rainfall and temperature trends in the study period was not consistent with the mean NDVI trends of wet and dry seasons. This result directly reflects the influence of rainfall and temperature was not considered as main drivers on the extensive change of vegetation cover of KSNP. Due to limited local meteorological stations, lack of advanced recording instruments, and skilled manpower leads to low accuracy of climate variables data. Therefore, the statistical analysis revealed precipitation and temperature did not consider as the main drivers of KSNP vegetation dynamics rather human-induced factors were the major actors for vegetation cover decline and degradation. Moreover, some dry season vegetation areas particularly along the Tekeze Riversides they are independent of annual rainfall as directly accessed water from the bank of the River. However, the seasonal variation/decreasing trends of NDVI in KSNP are more interlinked with increasing human drove activities such as the conversion of woody vegetation area to seasonal cultivation and bareland, loss of vegetation through dry season fire, firewood collection, and charcoal production than a change of precipitation and temperature. In line with our findings in dryland areas, change of vegetation cover in Nechsar National Park was due to degradation of vegetation via human driving deforestation [**41**]. Similarly, in the Kafta-humera district (around KSNP), Tigray region NDVI temporal reduction was due to human-induced changes in woodland vegetation [**40**]. The main human activities listed by the authors were the expansion of cropland, wood harvesting, and settlements. Moreover, a study in Africa Sahel also showed human-induced activities increase vegetation degradation [**125**]. However, the Ethiopian dryland vegetation productivity is dominantly controlled by the availability of moisture [**126**]; the low correlation between precipitation and NDVI is due to the decline of vegetation cover [**127**]. This is an indication of a little response of degraded woodland area to precipitation [**40**].

In contrast to the present study, reports witnessed that NDVI change (either increasing or decreasing) is driven by precipitation/temperature in arid and semi-arid areas of Africa [**128,129**] and in other countries (i.e Spain, Iraq and China) of the world [**94,130–133**]. NDVI controls the growth of vegetation conditions, temporal biomass accumulation, and changes [**134**]. The Spatiotemporal variation of NDVI was determined by the variation in the temporal distribution of precipitation [**135**]. On the other hand, the huge differences in the vegetation cover between dry and wet seasons were due to climatological and anthropogenic effects [**100**]. Moreover, in the Gojeb district, Ethiopia, and the three-north shelter forest of China vegetation degradation mainly influence by both human-induced factors and rainfall variability [**136,137**].

### Drivers of land use land cover change

The drivers of LULC change were both proximate and underlying [**109,115**]. For this discussion, more emphasis is given to the proximate drivers of LULC change. Based on the household survey, focus group discussion, and key informant interviews responded: settlement and agricultural expansion, illegal fire, and expansion of grazing, firewood collection, traditional gold mining, charcoal production, and the problem of land tenure as the main local communities recognized drivers of LULC change in KSNP. The main drivers were generated by a scarcity of resources in the moderate and high altitude areas of the region following resettlement from dense to less populated areas, low awareness in regard to environmental benefits of natural resources, lack of access to renewable energy sources (i.e solar energy and electric city) and high poverty.

According to the household survey, focus group discussion, and key informant interviewers’ resettlement schemes by the government and a few illegally were highly expanded in the study area between 1991 and 2003. From the interviewer result of local farmers around 71.7% were resettlers. Due to this resettlement program, new settlement communities’ namely: Tekeze, Maytemen, Maykeyh, Fre-Selam, Wohedet, and Mayweyni were established surrounding or nearest to KSNP. Similar studies showed resettlement program has brought significant LULC and massive devastating of the natural vegetation. For example, resettlement program 2003 in Chewaka district, Oromia region [**110**], in Gelda district of Amhara region between 1987 and 1990 [**87**], and in Esira district of South region [**138**] have resulted in a massive clearance of forest and depletion of much natural vegetation cover.

The majority of the sampled households (~94.7%) utilized wood and wood product for cooking. This activity leads to the dominant woodland destruction outside and inside KSNP. As stated by focus group discussion and key informant interview the woodland of the park is not only used as source energy but also used as generating alternative income by the local communities. This result is in line with Central RiftValley [**85**] and Malawi [**115**] they revealed fuelwood collection and charcoal making were the main drivers of LULC change. Resettlement program, expansion of farmland, human-induced fire, lack of land use plan, and unwise utilization were the top significant drivers of change [**110**]. Population growth, expansion of cultivation and settlement, fuelwood, and construction materials extraction from the forest was drivers of LULC change [**86**]. In Quirimbas National Park the drivers of LULC changes include; expansion of settlements and agriculture, forest resources trade, uncontrolled fires, infrastructures, exploitation of mining resources were reported [**20**].

The area has a great potential for non-renewable natural resources, traditional gold mining is a common activity and negatively changed the land cover of the study site. The activity of gold mining also leads to drift population from other areas towards the mine sites and increase settlements in the whole surrounding area of KSNP and even they constructed temporary structures as a place of residence inside the Park. According to vegetation surveyed of KSNP, all these listed activities destruct the natural vegetation via uprooting the plants’ root profile and a proximate cause for extinguishing illegal fire [**58**]. The emergence of gold mining activities coupled with an expansion of settlement in East Cameron had a negative impact on natural vegetation [**139**]. Furthermore, mining activities were clearly reported in Ghana as an important driver of LULC change and have a pronounced impact on the loss of vegetation [**118,140**].

Illegal fire is another proximate significant driver and directly related to LULC change in the studied site. The dry season fire of Kafta-sheraro national park was extinguished for a different purpose by three agents include (1) farmers (2) resin collectors and (3) traditional gold miners. The local community farmers prepared the land for cultivation through the destruction of woodlands by burning while resin collectors and traditional gold miners used fire for daily food preparation. All the agents destroyed the valuable natural resource of the park including wildlife migration outside the park. Detailed human-induced fire hazard related to LULC change report is limited in Ethiopia; however, studies undertaken from different corners of the world attest to the present result. A consistent study in hot spot tropical countries of South America (Mato Grosso-Brazil), East Africa (Serengeti ecological unit), and Central Africa (Cameroon) for 18 months period daily fire observation record (April 1992-December 1993) indicated a direct relation between the occurrence of fire and LULC change [**141**]. On the other hand, a study in central Spain between 1975 and 1990 of frequently burned areas or affected by fire showed the pine woodland covered decrease while shrubland encroachment turned to increase after five years of fire attack [**142**]. A similar report also indicated-human induced fire had a great contribution to increase destruction or loss of vegetation and this directly lead expanded of bareland areas [**140**].

The land tenure system was another significant driver of LULC change specifically in rural communities. As a result, the land administration problem is a great issue raised by the communities of the studied area. According to the responded households and focus group discussion the chairman of the Kebele representative gives the legality of the landowner to a user is selective (corruption) and this leads to create a gap and expansion of insecure land tenure in the district because having users landholding certificates could be expected to determine the land tenure security as stated by [**143**]. Such kinds of activities also open a door to expand illegal and unreasonable land invasion for cultivation. In the surrounding of KSNP communities surveyed households about 14% lacks legal land permit. Many reports confirmed the negative impact of insecure land administration on LULC dynamics [**65, 111**].

In addition to the local community pressure on natural resources, civil war and border conflict are additional factors in Ethiopia. According to the household, focus group discussion, and key informant interview response the Ethio-Eritrea war of 1998/99 which was dominantly destroyed socio-economic features of the Tigray region including the uncounted vegetation and wildlife resources in and around KSNP. During the war period, the military mechanized of the Eritrean troop crossed the park area, and destroyed the natural resources. This report is inconsistent with what happened in the Syria conflict (2011 to 2018) significantly changes the LULC from 2010 to 2018 and the conflict influenced local environmental resources [**144**]. Similarly, the civil war of Sri Lanka (1983 to 2009) and the post-war assessment of LULC change between 1993 and 2018 influenced forest reserves and protected areas of the country [**145**]. During the war period of Sierra Leones (civil war 1991-2002) caused LULC change of Kono district, forest cover structure and spatial extent was contracted [**146**].

### LULC change implication impact on park sustainability

Understanding and evaluating the spatial and temporal LULC change of natural vegetation in and outside protected areas and the interference of humans on these changes are significant. According to the classified image, change was detected in different land cover classes between 1988 and 2018 in KSNP and this indicated dominant woodland and riparian forest transformed into cultivation, grazing land and bareland. As a consequence the land surface of left with scattered vegetation and bare ground and this immediately showed how the natural vegetation overtime losses its plant biodiversity and limits the home range of the wildlife habitat. The changes of woodland drastically expanding into farmland and bareland in our study leading to encroachment of the elephant habitat and made a big challenge for their movement range.

According to our questionnaire assessment also confirmed the range of wildlife before 30 years were in everywhere of the park area. Recently, the suitable habitats for wildlife have shrunk and collapsed in specific areas of the park because agricultural expansion coupled with the extinguishing of illegal fire progressively increased from time to time. Secondly, the encroachment of livestock, firewood and charcoal collecter, and traditional gold miner leads to disturbance and displaced wildlife from the long existed habitat and shifted to new habitat patches. The increasing demand of the local communities’ for irrigated crop cultivation, livestock grazing, water for home and livestock consumption, and other resources utilization was an obstacle for wildlife movement and access to water especially, during the dry season. Such kinds of activities increased the conflict between elephants and humans. Moreover, the shift of riparian forest LULC class into farmland (irrigated area) around water points is the immediate cause for wildlife free movement and blocks water access. So, the wildlife is forced to migrate outside the park boundary even far distance to neighboring countries. In areas exposed by LULC change dramatically reduced the biodiversity [**1**]. Similar studies in Indonesia showed agricultural expansion caused continuous habitat quality decline coupled with a loss of biodiversity [**9**]. In a study in the Northern highlands of Ethiopia (1964 to 2014), LULC change was reported as an indication of plant and wildlife species loss [**86**]. A study in PAs of Mexico (1973 to 2015) indicated a reduction of temperate and tropical vegetation cover threatened the whole biodiversity of the site [**47**]. A recent study in Bale Mountain national park (1985 to 2017) revealed that the decreasing trends of grassland and forestland while increasing farmland LULC leads to increase habitat fragmentation and reduces in size and loss of available core area for the existing core dependent endemic wildlife species [**105**]. Furthermore, the expansion of agriculture had a threat to the general ecological integrity of elephant habitat and increased pressure for a competition of resources between humans and elephants [**43**]. The increase of bareland and decline of woody savanna in Tarangire national park was an indication of a threat for wildlife conservation [**101**].

## Conclusion

The results of three decades (1988-2018) period study have indicated extensive and significant LULC change was observed in KSNP. The degradation of woodland and riparian vegetation of the park resulting in the encroachment of shrub-bushland increases in sparse vegetation cover ground (bare land), cultivated land, and pasture land. The major depletion trend was observed from woodland to rainfed crop cultivation and from riparian vegetation to irrigated land. Moreover, bare land and cultivation enormously expanded from wet season to drier season. The result of NDVI analysis indicated the dense woodland and riverside vegetation decreased while non-vegetation increased between 1988 and 2018. Further, bare land was pronounced expansion from wet months to drier months.The gradual change in land use land cover of the park was mainly driven due to increasing human-induced pressure on the natural resources. The increasing trend of LULC change directly and negatively influences the wildlife habitats as the area is known home to African elephants and other large mammals. The major habitat change factors are the expansion of settlement and riverside/rainfed cultivation, human-induced fire, grazing, firewood collection, traditional gold mining, charcoal production, land administration problem, and natural resin collection which were degraded the wildlife habitat and limited their movement routes. Therefore, understanding the past and present LULC change drivers which are interlinked with the livelihood of the local communities is essential. Attention should be given to the sustainable conservation of park biodiversity by encouraging community participation in conservation practices, preparing awareness creation programs, and controlling all illegal activities practiced in and around KSNP. Further, this study is baseline information for setting effective land use planning and advanced management option of the park.

## Supporting information

**Appendix S1**: Detailed socio-economic study questionnaire form for households both in Tigrigna (local language) and English language.

## Acknowledgments

We greatly thank Ethiopian wildlife conservation authority (EWCA) for permiting this research site. We would also thank KSNP scouts and the local community of the districts for supporting field study and responding for questions and group discussion.

## Author contributions

Data collection, analysis, and writing original draft: Fitsum Temesgen; Supervision and editing: Bikila Warkineh, Alemayehu Hailemicael.

## References

1. Ellis E. Landuseand landcover change. In: Encyclopedia of Earth. Cleveland C.J. (eds). Environmental Information Coalition, National Council for Science and the Environment. 2006; http://www.eoearth.org/article/Landuselandcoverchange.

2. Garrard R, Kohler T, Price M.F, Byers AC, Sherpa AR, Maharjan G.R. Land Use and Land Cover Change in Sagarmatha National Park, a World Heritage Site in the Himalayas of Eastern Nepal. Mountain Research and Development. 2016; 36(3):299–310.

3. Rawat JS, Biswas V, Kumar M. Changes in land use/land cover using geospatial techniques: A casestudy of Ramnagar town area, district Nainital Uttarakhand, India. Egypt. J. Remote Sens. Space Science. 2013; 16:111–117.

4. Quentin FB, Jim C, Julia C, Carole H, Andrew S. Drivers of land use change, Final report: Matching opportunities to motivations, ESAI project 05116, Department of Sustainability and Environment and primary industries, Royal Melbourne Institute of Technology 2006.

5. Wondrade N, Dick ØB, Tveite H. GIS based mapping of land cover changes utilizing multi-temporal remotely sensed image data in Lake Hawassa Watershed, Ethiopia. Environmental Monitoring and Assessment. 2014; 186:1765–1780.

6. Singh RK, Singha M, Singh SK, Pal D, Tripathi N, Singh R. Land use/land cover change detection analysis using remote sensing and GIS of Dhanbad district, India. Eurasian Journal of Forest Science. 2018; 6(2):1–12.

7. Halmy MWA, Gessler PE, Hicke JA, Salem BB. Land use/land cover change detection and prediction in the north-western coastal desert of Egypt using Markov-CA. Appl. Geogr. 2015;63:101–112.

8. Turner MG, Ruscher CL. Change in landscape patterns in Georgia. USA. Land Ecology. 2004; 1(4):251–421.

9. Sharma R., Nehren U, Rahman SA, Meyer M, Rimal B, Seta G A, Baral H. Modeling Land Use and Land Cover Changes and their Effects on Biodiversity in Central Kalimantan, Indonesia. Land. 2018; 7(57):14p.

10. Butchart S. H. M., Walpole, M., Collen, B., Van Strien, A., Scharlemann, J. P. W., Almond, R. E. A., and Watson, R. Global biodiversity: Indicators of recent declines. Science. 2010; 328(5982):1164–1168.

11. Krauss J, Bommarco R, Guardiola M, Heikkinen R K, Helm A, Kuussaari M, Steffan D I. Habitat fragmentation causes immediate and time-delayed biodiversity loss at different trophic levels. Ecology Letters. 2010; 13:597–605.

12. Pimm SL, Raven P. Biodiversity: Extinction by numbers. Nature. 2000; 403:843–845.

13. Schmitz C, van Meijl H, Kyle P, Nelson GC, Fujimori S, Gurgel A, Havlik P, Heyhoe E. Land-use change trajectories up to 2050: Insights from a global agro-economic model comparison. Agric. Econ. 2014; 45:69–84.

14. Keesing F, Belden LK, Daszak P, Dobson A, Harvell CD, Holt RD, Ostfeld RS. Impacts of biodiversity on the emergence and transmission of infectious diseases. Nature. 2010; 468:647–652.

15. Worm B, Barbier EB, Beaumont N, Duffy JE, Folke C, Halpern BS, Watson R. The impact of biodiversity loss on ocean ecosystem services. Science. 2006; 314:787–790.

16. Tesfaw AT, Pfaff A, Kroner REG, Qin S, Medeiros R, Mascia MB. Land use land cover change shape the sustainability and impacts of protected areas. PNAS. 2018;6p. www.pnas.org/cgi/doi/10.1073/pnas.1716462115.

17. Butsic V, Baumann M, Shortland A, Walker S, Kuemmerle T. Conservation and conflict in the Democratic Republic of Congo: the impacts of warfare, mining, and protected areas on deforestation. Biological Conserv. 2015;191: 266–273.

18. Coetzee BWT, Gaston KJ, Chown SL. Local scale comparisons of biodiversity as a test for global protected area ecological performance: A meta-analysis. PLoS ONE. 2014; 9:e105824.

19. Geldmann J, Barnes M, Coad L, Craigie ID, Hockings M, Burgess ND. Effectiveness of terrestrial protected areas in reducing habitat loss and population declines. Biological Conservation. 2013;161:230–238.

20. Mucova SAR, Filho WL, Azeiteiro UM, Pereira MJ. Assessment of land use and land cover changes (1979-2017) and biodiversity and land management approach in Quirimbas National Park, Northern Mozambique, Africa. Global Ecology and Conservation. 2018; 16:e00447, 17p.

21. DeFries R, Hansen A, Newton AC, Hansen MC. Increasing isolation of protected areas in tropical forests over the past twenty years. Ecological Applications. 2005; 15:19–26.

22. Struhsaker TT, Struhsaker PJ, Siex KS. Conserving Africa’s rain forests: problems in protected areas and possible solutions. Biological Conservation. 2005;123:45–54.

23. Jones K.R, Venter O, Fuller RA, Allan JR, Maxwell SL, Negret PJ. et al. One-third of global protected land is under intense human pressure. Science. 2018; 360:788–791.

24. Sala OE, Chapin FS, Armesto JJ, Berlow E, Bloomfield J, Dirzo R, Wall DH. Global biodiversity scenarios for the year 2100, (New York). Science. 2000; 287:1770–1774.

25. Pereira HM, Leadley PW, Proenca V, Alkemade R, Scharlemann JPW, Fernandez-Manjarres J.F. et al. Scenarios for global biodiversity in the 21st century. Science. 2010; 330: 1496–1501.

26. Jones DA, Hansen AJ, Bly K, Doherty K, Verschuyl JP, Paugh JI, Story SJ. Monitoring land use and cover around parks: A conceptual approach. Remote Sensing of Environment. 2009; 113(7):1346–1356.

27. Hansen AJ, Defries R. Ecological mechanisms linking protected areas to surrounding lands. Ecological Applications. 2007; 17(4):974–988.

28. Beresford AE, Buchanan GM, Phalan B, Eshiamwata GW, Balmford A. et al. Correlates of long-term land-cover change and protected area performance at priority conservation sites in Africa. Environmental Conservation. 2018; 45(1):49–57.

29. Hellwig N, Walz A, Markovic D. Climatic and socioeconomic effects on land cover changes across Europe: Does protected area designation matter? PLoS ONE. 2019; 14(7):e0219374, 20p.

30. Bailey KM, McCleery RA, Binford MW, Christa Z. Land-cover change within and around protected areas in a biodiversity hotspot. Journal of Land Use Science. 2016; 11(2):154–176.

31. Hartter J, Southworth JD. Windling resources and fragmentation of landscapes around parks: Wetlands and forest patches around Kibale National Park, Uganda. Landscape Ecology. 2009; 24(5):643–656.

32. FAO. The state of the world’s land and water resources for food and agriculture (SOLAW)-managing systems at risk; Earthscan, London 2011;http://www.fao.org/docrep/03/i1688e/i1688e01. Accessed 10 may 2021. 281p.

33. Haines Y.R. Land use and biodiversity relationships. Land Use Policy. 2009; 26:178–86.

34. Lepers E, Lambin EF, Janetos AC, DeFries R, Achard F, Ramankutty N, Scholes RJ. A synthesis of information on rapid land-cover change for the period 1981-2000. Bioscience. 2005; 55:115–24.

35. Wilson EH, Sader SA. Detection of forest harvest type using multiple dates of Landsat TM imagery. Remote Sensing of Environment. 2002; 80:385–396.

36. Lambin EF, Geist HJ, Lepers E. Dynamics of land use/ cover change in tropical regions. Annual Review of Environment and Resources. 2003; 28:205–241.

37. Lemenih M, Kassa H, Kassie GT, Damtew A, Welday T. Resettlement and woodland management problems and options: A case study from North-Western Ethiopia. Land Degrad. Dev. 2014; 25:305–318.

38. Kassa T, Anton VR, Jean P, Simon VB, Jozef D, Kassa A. Spatial analysis of land cover changes in eastern tigray, Ethiopia from 1965 to 2007: Are there signs of a forest transition? Land Degrad Dev. 2014; doi:10.1002/ldr.2275.

39. Alemu B, Garedew E, Eshetu Z, Kassa H. Land Use and Land Cover Changes and Associated Driving Forces in North Western Lowlands of Ethiopia. International Research J. of Agricultural Science & Soil Science. 2015; 5(1):28–44.

40. Zewdie W, Csaplovics E. Identifying Categorical Land Use Transition and Land Degradation in North western Drylands of Ethiopia. Remote Sensing. 2016; 8(408):20p.

41. Fetene A, Hilker T, Yeshitela K, Prasse R, Cohen W, Yang Z. Detecting trends in Landuse and Landcover Change of Nech Sar National Park, Ethiopia. Environmental Management. 2015;11. DOI 10.1007/s00267-015-0603-0.

42. Nune S, Soromessa T, Teketay D. Land Use and Land Cover Change in the Bale Mountain Eco-Region of Ethiopia during 1985 to 2015. Land. 2016; 5(41): 22p.

43. Sintayehu DW, Kassaw M. Impact of land cover changes on elephant conservation in babile elephant sanctuary, Ethiopia. Biodiversity Int J. 2019; 3(2):65–71.

44. WoldeYohannes A, Cotter M, Kelboro G, Dessalegn W. Land Use and Land Cover Changes and their Effects on the Landscape of Abaya-Chamo Basin, Southern Ethiopia. Land. 2018; 7(2):17p.

45. Lu X, Zhou Y, Liu Y, LePage Y. The roles of protected areas in land use land cover change and the carbon cycle in the conterminous United States. Glob Change Biol. 2018; 24:617–630.

46. Duraisamy V, Bendapudi R, Jadhav A. Identifying hotspots in land use land cover change and the drivers in a semi-arid region of India. Environ Monit Assess. 2018; 190(535):21.

47. Sancheza Reyes UJ, Nino-Maldonado S, Barrientos-Lozano L, Trevino-Carreon J. Assessment of Land Use-Cover Changes and Successional Stages of Vegetation in the Natural Protected Area Altas Cumbres, Northeastern Mexico, Using Landsat Satellite Imagery. Remote Sensing. 2017; 9(712):33p.

48. Karakas CB, Cerit O, Kavak KS. Determination of land use land cover change and land use potentials of Sivas city and its surroundings using Geographical information systems and remote sensing. Procedia Earth and Planetary sciences. 2015; 15:454–461.

49. Tekle K, Hedlund L. Land cover changes between 1958 and 1986 in Kalu District, Southern Wello, Ethiopia. Mountain Research and Development. 2000; 20(1):42–51.

50. Karakus CB, Kavak KS, Cerit O. Determination of Variations in Land cover and Land use by Remote Sensing and Geographic Information Systems Around the City of Sivas (Turkey). Fresenius Environmental Bulletin. 2014; 23(3):667–677.

51. Rawat JS, Kumar M. Monitoring land use/cover change using remote sensing and GIS techniques: A case study of Hawalbagh block, district Almora, Uttarak hand, India. The Egyptian Journal of Remote Sensing and Space Sciences. 2015; 18:77–84.

52. Diallo Y, Hu G, Wen X. Applications of Remote Sensing in Land Use/Land Cover Change Detection in Puer and Simao Counties, Yunnan Province. Journal of American Science. 2009; 5(4):157–166.

53. Lambin EF, Meyfroidt P. Land use transitions: Socio-ecological feedback versus socio-economic change. Land Use Policy. 2010;27:108–118.

54. Kamwi JM, Cho MA, Kaetsch C, Manda SO, Graz FP, Chirwa PW. Assessing the Spatial Drivers of Land Use and Land Cover Change in the Protected and Communal Areas of the Zambezi Region, Namibia. Land. 2018; 7(131):13p.

55. Getahun K, Poesen J, Rompaey AV. Impacts of Resettlement Programs on Deforestation of Moist Evergreen Afromontane Forests in Southwest Ethiopia. Mountain Research and Development. 2017; 37(4): 474–486.

56. Wulder MA, White JC, Cranny M, Hall RJ, Luther JE, Beaudoin A. et al. Monitoring Canada’s forests. Part 1: completion of the EOSD land cover project. Canadian Journal of Remote Sensing. 2008; 34(6):549–562.

57. Cohen WB, Goward S. Landsat’s role in ecological applications of Remote Sensing. Bioscience. 2004, 54:535–545.

58. Temesgen F, Warkineh B. Woody Species Structureand Regeneration Status in Kafta Sheraro National Park Dry Forest, Tigray Region, Ethiopia. International Journal of Forestry Research. 2020;Volume 2020: Article ID 4597456, 22p.

59. Dejene T, Lemenih M, Bongers F. Manage or convert boswellia woodlands? Can frankincense production pay off? Journal of Arid Environment. 2013; 89:77–83.

60. Ethiopian National Meteorology Agency of Rainfall and Temperature data (1996-2016), Addis Ababa, Ethiopia 2018.

61. Cochran WG. Sampling techniques, 3^rd^ ed. John Willey and Sons, New York. 1977.

65. Ewunetu A, Simane B, Teferi E, Zaitchik BF. Land Cover Change in the Blue Nile River Headwaters: Farmers’ Perceptions, Pressures, and Satellite-Based Mapping. Land. 2021; 10(68):1–25.

66. USAID. Conducting key informants’ interviews. Center for development information and evaluation 1996; 1–4p.

67. Coppin P, Jonckheere I, Nackaerts K, Muys B, Lambin E. Digital change detection methods in ecosystem monitoring: A review. International journal of remote sensing. 2004; 25(9):1565–1596.

68. Rani N, Mandla VR, Singh T. Evaluation of atmospheric corrections on hyperspectral data with special reference to mineral mapping. Geoscience Frontiers. 2017; 8(4):797–808.

69. Hansen MC, Loveland, T R. A review of large area monitoring of land cover change using Landsat data. Remote Sensing of Environment. 2012; 122:66–74.

70. Hilker T, Lyapustin AI, Tucker CJ, Sellers PJ, Hall FG, Wang Y. Remote sensing of tropical ecosystems: atmospheric correction and cloud masking matter. Rem. Sens. Environ. 2012; 127: 370–384.

71. Chander G, Markham BL, Helder DL. Summary of current radiometric calibration coefficients for Landsat MSS, TM, ETM^+^, and EO-1 ALI sensors. Remote Sensing Environment. 2009; 113:893–903.

72. Rogan J, Chen D. “Remote Sensing Technology for Mapping and Monitoring Land Cover and Land-use Change.” Progress in Planning. 2004; 61:301–325.

73. Abyot Y, Berhanu G, Solomon A, Ferede Z. Forest cover change detection using remote sensing and GIS in Banja district, Amhara region, Ethiopia. International Journal of Environmental Monitoring & Analysis. 2014; 2:354–360.

74. Manandhar R, Odeh IOA, Ancev T. Improving the Accuracy of Land Use and Land Cover Classification of Landsat Data using Post-classification Enhancement. Remote Sensing. 2009; 1:330–344.

75. Yuan F, Sawaya KE, Loeffelholz BC, Bauer ME. Land cover classification and change analysis of the Twin cities (Minnesota) Metropolitan Area by multi-temporal Landsat remote sensing. Remote Sensing Envir. 2005; 98:317–328.

76. Wu W, Shao G. “Optimal Combinations of Data, Classifiers, and Sampling Methods for Accurate Characterization of Deforestation. Canadian Journal of Remote Sensing. 2002; 28(4):601–609.

77. Othow OO, Gebre SL, Gemeda D.O. Analyzing the Rate of Land Use Land Cover Change and Determining the Causes of Forest Cover Change in Gog District, Gambella, Ethiopia. Journal Remote Sensing and GIS. 2017; 6:219.

78. Congalton R, Green K. Assessing the accuracy of remotely sensed data: Principles and practices. 2^nd^ ed. Taylor and Francis Group, Boca Raton, USA. 2009; 168p.

79. Keshtkar H, Voigt W, Alizadeh E. Land-cover classification and analysis of change using machine learning classifiers and multi-temporal remote sensing imagery. Arabian Journal of Geo-sciences. 2017; 10(6).

80. Campbell JB. Introduction to remote sensing. 2^nd^ ed. London: Taylor and Francis 1996.

81. Smits PC, Dellepiane SG, Schowengerdt RA. Quality assessment of image classification algorithms for land-cover mapping: a review and proposal for a cost-based approach. Inter. J. Remote Sens. 1999; 20:1461–86.

82. Lillesand T, Kiefer R. Chipman J. Remote Sensing and Image Interpretation. 6^th^ ed. John Wiley and Sons, New York, USA. 2008; 756p.

83. Sim J, Wright CC. The Kappa Statistic in Reliability Studies: Use, interpretation, and sample size Requirements. Phys. Ther. 2005; 85:257–68.

84. Mahmud A, Achide A S. Analysis of Land Use/Land Cover Changes to Monitor Urban Sprawl in Keffi-Nigeria. Environmental Research Journal. 2012; 6:129–134.

85. Bekele B, Wu W, Yirsaw E. Drivers of Land Use-Land Cover Changes in the Central Rift Valley of Ethiopia. Sains Malaysiana. 2019; 48(7):333–1345.

86. Asmame B, Abegaz A. Land use/land cover changes and their environmental implications in the Gelana sub-watershed of Northern highlands of Ethiopia. Environ Syst Res. 2017; 6(7):12p.

87. Esa E, Assen M. Land use/cover dynamics and its drivers in Gelda catchment, Lake Tana watershed, Ethiopia. Environmental Systems Research. 2017; 6(4):1–13.

88. Alawamy JSSK, Balasundram A, Hanif H, Sung CT. Detecting and Analyzing Land Use and Land Cover Changes in the Region of Al-Jabal Al-Akhdar, Libya Using Time-Series Landsat Data from 1985 to 2017. Sustainability. 2020; 12(4490):24p.

89. Worku T, Tripathi S K, Khar D. Analyses of land use and land-cover change dynamics using GIS and remote sensing during 1984 and 2015 in the Beressa Watershed Northern Central Highland of Ethiopia. Model. Earth Syst. Environ. 2016; 2(168):1–12.

90. MeeraGandhi G, Parthiban S, Thummalu N, Christy A. Ndvi: Vegetation change detection using remote sensing and gis -A case study of Vellore District. Procedia Computer Science. 2015; 57:1199–1210.

91. Meneses-Tovar CL. NDVI as indicator of degradation. Unasylva. 2011; 238(62):40–46.

92. Schnur MT, Hongjie X, Wang X. Estimating root zone soil moisture at distant sites using MODIS NDVI and EVI in a semiarid region of southwestern USA. Ecol. Inform. 2010; 5:400–409.

93. Naif SS, Mahmood DA, Al-Jiboori MH. Seasonal normalized difference vegetation index responses. Open Agriculture. 2020; 5:631–637.

94. Bilgili BCO, Satir V, Muftuoglu M, Ozyavuz A. Simplified Method for the Determination and Monitoring of Green Areas in Urban Parks Using Multispectral Vegetation Indices. J Environ Prot Ecol. 2014;15(3):1059.

95. Musa L, Peters K, Ahmed M. On farm characterization of Butana and Kenana cattle breed production systems in Sudan. Livest. Res. Rural Dev. 2006; 18:1277–1288.

96. Lesschen JP, Verburg PH, Staal SJ. Statistical Methods for Analysing the Spatial Dimension of Changes in Land Use and Farming Systems; International Livestock Research Institute: Nairobi, Wageningen. 2005;80p.

97. R-Development Core Team. R: A Language and Environment for Statistical Computing, R-Foundation for Statistical Computing, Vienna, Austria 2019.

98. IBM Statistical Package for Social Sciences (SPSS) Statistics for Windows, Version 26.0. Armonk, New York:IBM Corp 2019.

99. Mesfin D, Simane B, Belay A, Recha JW, Taddese H. Woodland Cover Change in the Central Rift Valley of Ethiopia. Forests. 2020; 11(916):16p.

100. Al-Saady Y, Merkel B, Al-Tawash B, Al-Suhail Q. Land use and land cover (LULC) mapping and change detection in the Little Zab River Basin (LZRB), Kurdistan Region, NE Iraq and NW Iran. Freiberg Online Geoscience. 2015; 43:32p

101. Mtui DT, Lepczyk CA, Chen Q, Miura T, Cox LJ. Assessing multi-decadal land-cover -land-use change in two wildlife protected areas in Tanzania using Landsat imagery. PLoS ONE. 2017; 12(9):e0185468, 20p.

102. Liping C, Yujun S, Saeed S. Monitoring and predicting land use and land cover changes using remote sensing and GIS techniques DA case study of a hilly area, Jiangle, China. PloS ONE. 2018; 13(7): e0200493.

103. Msigwa A, Komakech HC, Verbeiren B, Salvadore E, Hessels T, Weerasinghe I, Griensven A. Accounting for Seasonal Land Use Dynamics to Improve Estimation of Agricultural Irrigation Water Withdrawals. Water. 2019; 11(2471):20p.

104. Liu C, Frazier P, Kumar L. Comparative assessment of the measures of thematic classification accuracy. Remote Sens. Environ. 2007; 107:606–616.

105. Muhammed A, Elias E. Class and landscape level habitat fragmentation analysis in the Bale mountains national park, southeastern Ethiopia. Heliyon. 2021;7:e07642.

106. Solomon B, Amsalu A, Abebe E. Land Use and Land Cover Changes in Awash National Park, Ethiopia: Impact of Decentralization on the Use and Management of Resources. Open Journal of Ecology, 2014;4: 950–960.

107. Garedew E, Sandewall M, Soderberg U, Campbell B. Land-use and land-cover dynamics in the central rift valley of Ethiopia. Environmental Management. 2009; 44(4):683–694.

108. Berihun ML, Tsunekawac A, Haregeweyn N, Mesheshae DT, Adgoe E, Tsuboc M. et al. Exploring land use/land cover changes, drivers and their implications in contrasting agro-ecological environments of Ethiopia. Land Use Policy. 2019; 87:104052.15p.

109. Bewket W, Abebe S. Land-use and land-cover change and its environmental implications in a tropical highland watershed, Ethiopia. International Journal of Environmental Studies. 2013; 70(1):126–139.

110. Abera A, Yirgu T, Uncha A. Impact of resettlement scheme on vegetation cover and its implications on conservation in Chewaka district of Ethiopia. Environmental System Resources. 2020; 9:2:17.

111. Gedefaw AA, Atzberger C, Bauer T, Agegnehu SK, Mansberger R. Analysis of Land Cover Change Detection in Gozamin District, Ethiopia: From Remote Sensing and DPSIR Perspectives. Sustainability. 2020; 12(4534):1–25.

112. Erdogan HE, Pellikka PKE, Clark B. Modelling the impact of land-cover change on potential soil loss in the Taita Hills, Kenya, between 1987 and 2003 using remote-sensing and geospatial data. Int. J. Remote Sens. 2011; 32:5919–5945.

113. Maeda EE, Clark BJF, Pellikka P, Siljander M. Modelling agricultural expansion in Kenya’s Eastern Arc Mountains biodiversity hotspot. Agric. Syst. 2010; 103:609–620.

114. Egeru A, Wasonga O, Kyagulanyi J, Majaliwa GJM, MacOpiyo L, Mburu J. Spatio-temporal dynamics of forage and land cover changes in Karamoja sub-region, Uganda. Pastoralism: Research, Policy and Practice. 2014; 4(6):1–21.

115. Munthali MG, Davis N, Adeola AM, Botai JO, Kamwi JM, Chisale HLW. Local perception of drivers of land-use and land-cover change dynamics across Dedza district, Central Malawi Region. Sustainability. 2019; 11(832):25p.

116. Schurmanna A, Kleemanna J, Fursta C, Teucherb M. Assessing the relationship between land tenure issues and land cover changes around the Arabuko Sokoke Forest in Kenya. Land Use Policy. 2020; 95(104625):1–13.

117. How Jin Aik D, Ismail MH, Muharam FM, Alias MA. Evaluating the impacts of land use/land cover changes across topography against land surface temperature in Cameron Highlands. PLoS One. 2021; 16(5):e0252111, 1-26.

118. Garai D, Narayana AC. Land use/land cover changes in the mining area of Godavari coal fields of Southern India. The Egyptian Journal of Remote Sensing and Space Sciences. 2018; 21:375–381.

119. Pham TT, Nguyen M, The-Duoc Tham HTN, Truong TNK, Lam-Dao N, Nguyen-Huy T. Specifying the relationship between land use/land cover change and dryness in central Vietnam from 2000 to 2019 using Google Earth Engine. Journal of Applied Remote Sensing. 2021; 15(2):1–21.

120. Hassan Z, Shabbir R, Ahmad SS, Malik AH, Aziz N, Butt A, Erum S. Dynamics of land use and land cover change (LULCC) using geospatial techniques: a case study of Islamabad Pakistan. Springer Plus. 2016; 5(812):1–11.

121. Wingate VR, Phinn, SR, Kuhn N, Bloemertz L, Dhanjal-Adams KL. Mapping Decadal Land Cover Changes in the Woodlands of North Eastern Namibia from 1975 to 2014 Using the Landsat Satellite Archived Data. Remote Sensing. 2016; 8(681):20p.

122. Maina J, Wandiga S, Gyampoh B, Charles KKG. Assessment of Land Use and Land Cover Change Using GIS and Remote Sensing: A Case Study of Kieni, Central Kenya. J Remote Sens GIS. 2020; 9:270.

123. Amri R, Zribi M, Lili-Chabaane Z, Duchemin B, Gruhier C, Chehbouni A. Analysis of vegetation behavior in a North African semi-arid region, using SPOT-vegetation NDVI data. Remote Sens. 2011; 3, 2568–2590.

124. Ferreira LG, Yoshioka H, Huete A, Sano EE. Seasonal landscape and spectral vegetation index dynamics in the Brazilian Cerrado: An analysis within the Large-Scale Biosphere-Atmosphere Experiment in Amazônia (LBA). Remote Sens. Environ. 2003; 87, 534–550.

125. Evans J, Geerken R. Discrimination between climate and human-induced dryland degradation, Journal of Arid Environments, 2004; 57:535–554.

126. Hailu BT, Maeda EE, Heiskanen J, Pellikka P. Reconstructing pre-agricultural expansion vegetation cover of Ethiopia. Appl. Geogr. 2015; 62:357–365.

127. Li J, Lewis J, Rowland J, Tappan G, Tieszen LL. Evaluation of land performance in Senegal using multi-temporal NDVI and rainfall series. J. Arid Environ. 2004; 59:463–480.

128. Ghebrezgabher MG, Yang T, Yang X, Sereke TE. Assessment of NDVI variations in responses to climate change in the Horn of Africa. The Egyptian Journal of Remote Sensing and Space Sciences. 2020; 23:249–261.

129. Martiny N, Camberlin P, Richard Y, Philippon N. Compared regimes of NDVI and rainfall in semi-arid regions of Africa. International Journal of Remote Sensing. 2006; 27(23):5201–5223.

130. Chu D, Lu L, Zhang T. Sensitivity of Normalized Difference Vegetation Index (NDVI) to Seasonal and Interannual Climate Conditions in the Lhasa Area, Tibetan Plateau, China. Arctic, Antarctic, and Alpine Research. 2007; 39(4):635–641.

131. Fensholt R, Langanke T, Rasmussen K, Reenberg A, Prince SD, Tucker C. et al. “Greenness in semi-arid areas across the globe 1981-2007-an Earth Observing Satellite based analysis of trends and drivers,” Remote Sensing Environment. 2012; 121:144–158.

132. Pei Z, Fang S, Yang W, Wang L, Wu M, Zhang Q, Han W, Khoi DN. The Relationship between NDVI and Climate Factors at Different Monthly Time Scales: A Case Study of Grasslands in Inner Mongolia, China (1982-2015). Sustainability. 2019; 11(7243):1–17.

133. Sanz E, Saa-Requejo A, Díaz-Ambrona CH, Ruiz-Ramos M, Rodríguez A, Iglesias P. et al. Normalized Difference Vegetation Index Temporal Responses to Temperature and Precipitation in Arid Rangelands. Remote Sens. 2021; 13(840):1–24.

134. Zhao F, Xu B, Yang X, Jin Y, Li J, Xia L, Chen S, Ma H. Remote sensing estimates of grassland aboveground biomass based on MODIS Net Primary Productivity (NPP): A case study in the Xilingol Grassland of northern China. Remote Sens. 2014;6:5368–5386.

135. Xia J, Liu S, Liang S, Chen Y, Xu W, Yuan W. Spatio-temporal patterns and climate variables controlling of biomass carbon stock of global grassland ecosystems from 1982 to 2006. Remote Sens. 2014; 6: 1783–1802.

136. Huang S, Kong J. “Assessing land degradation dynamics and distinguishing human-induced changes from climate factors in the three-north shelter forest region of China,” ISPRS International Journal of Geo-Information. 2016; 5(9):158.

137. Dagnachew M, Kebede A, Moges A, Abebe A. Effects of Climate Variability on Normalized Difference Vegetation Index (NDVI) in the Gojeb River Catchment, Ethiopia. Advances in Meteorology. 2020; Article ID 8263246, 1–16.

138. Mamude M, Melka GA, Genet W. Geospatial Techniques Based Analysis on the Impact of Resettlement on Land Use and Land Cover Change in Esira District, Dawuro Zone, Ethiopia. Ghana Journal of Geography. 2021; 13(1): 203–220.

139. Kamga MA, Fils SCN, Ayodele MO, Olatubara CO, Nzali S, Adenikinju A, Khalifa M. Evaluation of land use/land cover changes due to gold mining activities from 1987 to 2017 using landsat imagery, East Cameroon. GeoJournal. 2020; 85:1097–1114.

140. Prosper LB, Guan Q, Cheng D. Exploring Land use and Land cover change in the mining areas of Wa East District, Ghana using Satellite Imagery. Open Geosci. 2015; 1:618–626.

141. Eva H, Lambin EF. Fires and land-cover change in the tropics: a remote sensing analysis at the landscape scale. Journal of Biogeography. 2000;27:765–776.

142. Viedma O, Moreno Jm, Rieiro I. Interaction between land use/land cover change,forest fires and land scape structure in Sierrra de Gredos (central spain). Environmental Conservation. 2006; 33(3):212–222.

143. Tsegaye A, Adgo EG, Selassie Y. Impact of Land Certification on Sustainable Land Resource Management in Dryland Areas of Eastern Amhara Region, Ethiopia. J. Agric. Sci. 2012; 4:261–268.

144. Mohamed Al, Anders J, Schneider C. Monitoring of Changes in Land Use/Land Cover in Syria from 2010 to 2018 Using Multitemporal Landsat Imagery and GIS. Land. 2020; 9(226):1–30.

145. Rathnayake CWM, Jones S, Soto-Berelov M. Mapping Land Cover Change over a 25 Year Period (1993-2018) in Sri Lanka Using Landsat Time-Series. Land. 2020; 9(27):1–19.

146. Wilson SA, Wilson C O. Modelling the impacts of civil war on land use and land cover change within Kono District, Sierra Leone: A socio-geospatial approach. Geocarto International. 2012; 1–26.

